# Chemically modified CRISPR enzymes for multi-organ genome editing *in vivo*

**DOI:** 10.64898/2026.09.14.751351

**Authors:** Christopher M. Baehr, Alzbeta Ressnerova, Min Kang, Sheng Zhao, Beatrice Le, Rania F. Haddad, Kunica Asija, Mark Damante, Rohit Sharma, Christy George, Heidi Huang, Dror Assa, Eric A. Noel, Brigette Manohar, P. Anthony Otero, Meika Travis, Kayla Saikaly, Sophie Hu, Victor S. Van Laar, Russell R. Lonser, John G. Flannery, Niren Murthy, Ross C. Wilson

## Abstract

Delivery remains the main obstacle to the development of *in vivo* genome editing therapies. CRISPR ribonucleoproteins confer high editing activity with transient exposure but lack intrinsic cell entry and targeting. Here we introduce PERCEPT, a delivery platform featuring reversible, covalent modification of CRISPR enzymes. PERCEPT enables modular installation of shielding polymers, amphiphilic delivery peptides and targeting ligands, allowing traceless cytosolic release of the native editor. A formulation incorporating an amphiphilic delivery peptide and the neuron-targeting ligand TET1 edited approximately 56% of striatal volume and 78% of neurons within edited regions after local striatal administration. In the R6/2 Huntington’s disease model, PERCEPT mediated targeted editing of the mutant human *HTT* transgene, reducing mutant huntingtin aggregate burden and shifting local transcriptional programs away from inflammatory and injury-associated states. Tissue-adapted formulations edited 53% of Müller glia following intravitreal delivery, increased skeletal muscle reporter fluorescence tenfold versus unconjugated control following intramuscular injection, and enabled lung airway epithelial cell that persisted for three months following intranasal delivery. Rapid screening revealed that local anatomical and cellular barriers require distinct surface display for optimal editing. These results establish reversible chemical modification as a general strategy for adapting CRISPR enzyme delivery across multiple target tissues *in vivo*.

## Introduction

Delivery remains the main obstacle to the development of *in vivo* genome editing therapies^1,2^. Genome editors must traverse extracellular space, access disease-relevant cells, and remain transient to limit risks of genotoxicity and immunogenicity. CRISPR-Cas enzymes delivered as pre-formed ribonucleoprotein (RNP) enzymes provide transient exposure^3^, but efficient tissue access and cell targeting have been challenging to date. Viral vectors can provide tropism, but constitutive expression of CRISPR machinery burdens the desired one-time edit with the persisting liabilities of off-target editing, vector integration, and immune activation^4–6^. Lipid nanoparticles (LNPs) have demonstrated clinical delivery of genome editing machinery for liver editing^7^. However, delivery to solid extrahepatic tissue remains challenging^8,9^. Extracellular transport through brain parenchyma or lung mucus depends heavily on small particle size and favorable surface properties^10–12^. Established preclinical RNP delivery systems, including engineered virus-like particles (VLPs)^13–15^, extracellular vesicles^16^, inorganic nanoparticles^17^, RNP-CPP fusions^18^, nanocapsules^19^, and LNP formulations optimized for RNPs^20^, demonstrate activity in selected settings. However, these approaches generally do not provide reversible, modular, on-demand surface functionalization of the genome-editing cargo itself. As a result, these platforms offer limited flexibility for rapid tuning of RNP formulations for delivery to diverse tissues. Furthermore, many of these RNP-centric approaches eliminate a key advantage inherent to RNP enzymes: they are small biologics with a diameter below 20 nm, which is optimal for tissue distribution^21^ and compares favorably to the ≥100 nm diameter typical of CRISPR-bearing LNPs and VLPs^10–12^. An idealized RNP delivery platform would retain the enzyme’s small particle size while offering convenient optimization of surface properties.

We previously developed peptide-enabled RNP CRISPR (PERC) for *ex vivo* engineering of therapeutically-relevant cells that have been challenging to transduce, including T cells and hematopoietic stem cells^22,23^. PERC relies on engineered, virus-derived cell penetrating and endosomolytic fusion peptides (e.g. A5K or P55) that can be admixed with CRISPR RNPs, mediating efficient and minimally-toxic genome editing in some target cell types. However, the admixed PERC complexes are not readily compatible with the incorporation of diverse cell-targeting moieties, which would ideally be covalently associated with RNP cargo.

Here, we introduce PERCEPT (PERC engineered via polymeric tethering) an *in vivo* CRISPR delivery platform featuring a reversible, traceless linker that covalently couples chemical and biochemical moieties to RNP enzymes, decorating CRISPR-Cas9 nuclease (henceforth Cas9) and adenine base editors (ABE) for activity and cell specific targeting in multiple tissues. Building on our previous work^24^, the linker covalently reacts with primary amines (e.g., lysine side chains, of which Cas9 has >100 exposed to solvent), and incorporates a reduction-sensitive cleavage site that scarlessly restores native enzyme structure and function. The linker comprises polyethylene glycol (PEG) terminating in a distal bioorthogonal click handle that facilitates convenient, rapid, modular, and highly efficient conjugation to endosomolytic peptides, cell targeting moieties, and/or shielding polymers^25^. Together, these features comprise PERCEPT, a reversible, modular surface-programming platform for delivery of CRISPR RNPs whose delivery performance is tuned by tissue-specific chemistry.

Using PERCEPT, we show that programmable RNP surface chemistry can separately influence distribution, cellular uptake, and cell-type engagement *in vivo*. In murine striatum after convection-enhanced delivery, we observed that supplemental PEGylation improves distribution, an endosomolytic peptide enhances uptake and editing, and the neuron-targeting ligand TET1 increases neuronal enrichment. The same platform supports adenine base editor delivery as well as disease-relevant editing in an aggressive model of Huntington’s disease, reducing pathological aggregates and rescuing a disease-associated transcriptomic profile. Beyond the CNS, related – yet chemically distinct – formulations enabled editing in the retina, skeletal muscle, and airway epithelium, establishing PERCEPT as a versatile non-viral delivery platform for genome editing of multiple organs *in vivo*.

## RESULTS

### PERCEPT enables reversible programming of Cas9 RNP surface properties

We sought to develop a chemically defined Cas9 RNP delivery architecture that would retain the small particle size of the native RNP while incorporating functions required for intracellular delivery and cell targeting, with sufficient stability for formulation and handling. Our previously-reported PERC platform established a strategy for promoting cellular entry of Cas9 RNPs through admixture of RNP & delivery peptide^22,23^, while our prior work developing DEC-PEG demonstrated that polymers could be reversibly tethered to Cas9 through a self-immolative linker, allowing traceless recovery of the native protein after reduction mediated by free glutathione found in the cytosol^24^. Building on these advances, we designed PERCEPT to incorporate a bioorthogonal click handle at the distal PEG terminus, enabling modular conjugation of targeting and delivery ligands onto linker-modified RNPs. As an initial application, we focused on neuronal targeting using TET1 (henceforth T1), a peptide ligand previously shown to promote neuronal uptake^26^ (see results below). Together, these elements create a compact, chemically defined RNP architecture in which reversible protein modification, PEG shielding, and click-mediated installation of functional ligands can be independently incorporated (Fig. 1a).

**Figure 1.**
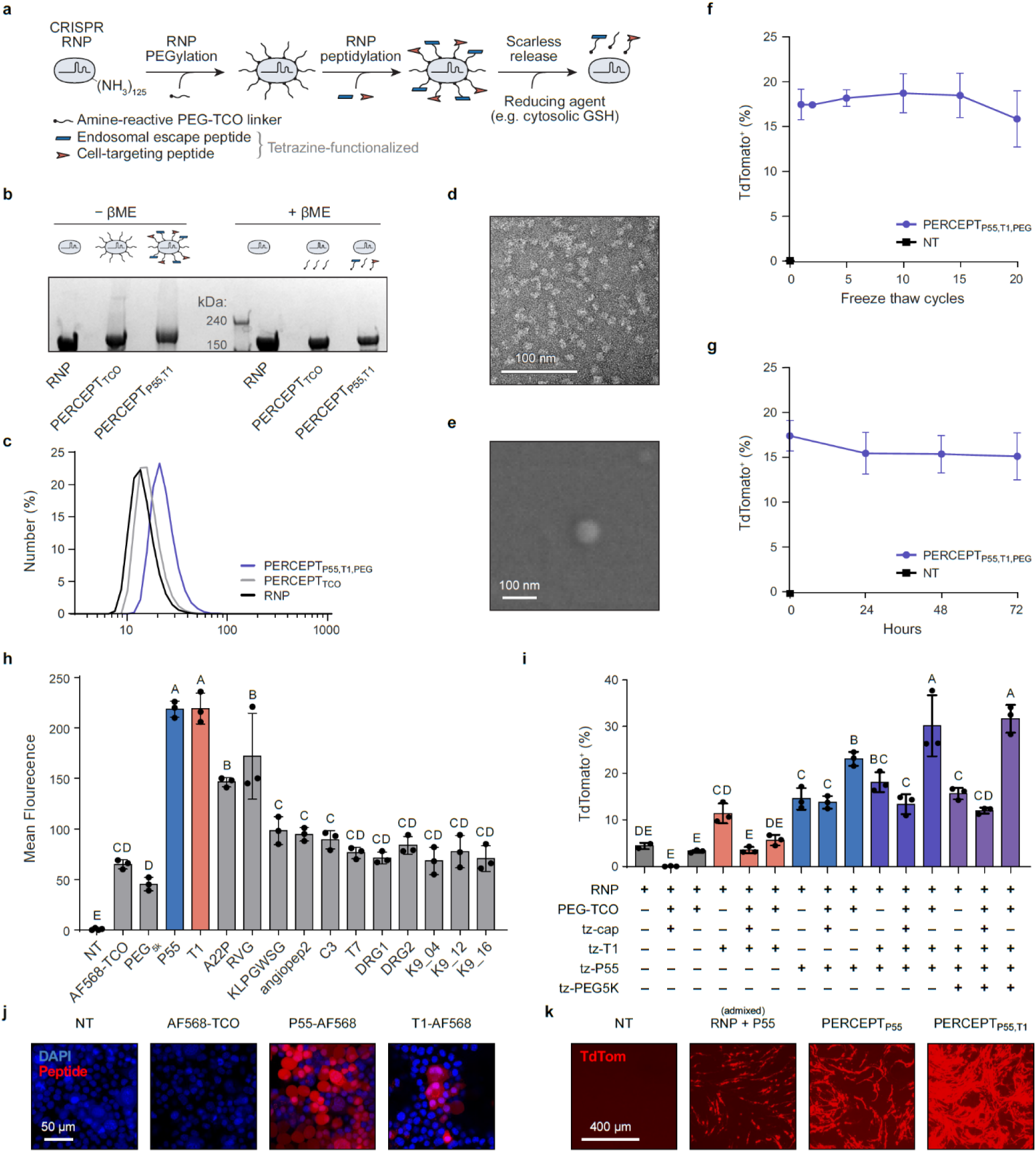
PERCEPT enables reversible, aqueous surface programming of Cas9 RNPs. a, Schematic of PERCEPT functionalization: aqueous installation of a reduction-sensitive PEG carbamate linker on accessible Cas9 RNP amines enables modular trans-cyclooctene–tetrazine ligation of delivery peptides, targeting ligands or shielding polymers; reduction triggers linker self-immolation and native RNP release. **b,** Denaturing gel analysis of unmodified, linker-modified and peptide-conjugated RNPs ± β-mercaptoethanol (βME). Modified species show reduced mobility without βME and collapse toward the native Cas9 band with βME, supporting reduction-sensitive loss of modification. **c,** RNP hydrodynamic diameter after sequential surface modification, measured by dynamic light scattering. **d,e,** Transmission electron microscopy of uranyl acetate-stained (d) and scanning electron microscopy of carbon-sputtered (e) PERCEPT RNPs. **f,g,** PERCEPT (P55, T1 and 5-kDa PEG) editing activity after indicated freeze–thaw cycles (f) or room-temperature incubation times (g), measured as percentage tdTomato-positive cells. **h,** Flow cytometry of cell-associated AF568 fluorescence in Neuro2A cells treated with indicated fluorophore conjugates or controls. **i,** Flow cytometry of tdTomato activation in Ai9-derived neural progenitor cells comparing unconjugated, admixed and covalently conjugated RNPs. Matrix, inclusion (+) or omission (−) of PEG-TCO linker, tetrazine cap, T1, P55 and 5-kDa PEG. **j,k,** Representative fluorescence microscopy of Neuro2A treatments showing DAPI and cell-associated AF568-labeled peptide (j), and Ai9-derived NPCs showing tdTomato activation after RNP treatments (k). For h,i, three replicate wells per sample (96-well plates). Letters denote compact letter displays from ANOVA with Tukey’s test; shared letters indicate no significant difference. Scale bars, 100 nm (d,e), 50 μm (j), 400 μm (k).

We first evaluated whether the PERCEPT linker could modify Cas9 RNPs under aqueous conditions while preserving the ability to release native protein after reduction. Native SpCas9 RNP, linker-bearing RNP, and peptide-conjugated RNP were evaluated via denaturing SDS-PAGE under reducing or non-reducing conditions. Under non-reducing conditions, the migration of linker-and peptide-modified samples was increasingly slowed as compared to native RNP, consistent with the two progressive steps of covalent surface modification. Under reducing conditions, the modified species collapsed to match the native SpCas9 band, indicating redox-triggered linker self-immolation and release of the native RNP cargo, as we expected based on our previous work^24^ (Fig. 1b; Extended Data Fig. 1).

We next characterized how sequential surface modification affected RNP size and morphology. Dynamic light scattering showed a stepwise increase in hydrodynamic diameter, from ∼12 nm for native SpCas9 RNP to ∼15 nm after linker installation and ∼22 nm after peptide conjugation. Negative-stain transmission electron microscopy corroborated this trend and showed predominantly compact particles within the same size range. Scanning electron microscopy further supported the compact morphology of PERCEPT-formulated RNPs. Nanogold labelling visualized linker-modified RNP valency (Extended Data Fig. 8). RNP activity remained active following freeze-thaw challenge and room-temperature incubation, demonstrating robustness during formulation handling. Thus, the chemical biology steps that generate PERCEPT formulations marginally increased RNP particle size (as expected) and maintained a stable ≤25 nm profile anticipated to promote efficient tissue distribution (Fig. 1c–g).

### Ligand screening identifies T1 as a neuron-associated targeting ligand

To promote CNS editing applications, we sought to identify a neuron-targeting ligand that could be incorporated into PERCEPT formulations. Tetrazine-modified candidate peptides identified from prior studies (P55^23^, T1^26^, A22p^18,27^, RVG^28^, KLPGWSG^29^, Angiopep2^30^, C3^31^, T7^32^, DRG1^33^, DRG2^33^) as well as a novel trio of evolved AAV-derived cyclic peptides (K9_04, K9_12 and K9_16^34^) were pre-complexed with TCO-AF568 to generate fluorescent peptide conjugates and applied to neuronal cells. Cell-associated AF568 fluorescence was quantified by flow cytometry then normalized to AF568-TCO alone, simultaneously fluorescence in adherent cells was visualized via confocal scanning laser microscopy (Fig. 1h,j).

P55, an amphiphilic HA2-TAT fusion peptide that promotes endosomal escape in a diversity of cell types as previously reported^23^, produced strong cell-associated fluorescence, reaching 3.4-fold over AF568-TCO. Among neuron-targeting candidates, T1 produced a comparable signal, also 3.4-fold over AF568-TCO, and was statistically grouped as one of the highest-performing ligands (along with P55). RVG and A22p produced intermediate signal, whereas the remaining candidates ranged from approximately baseline to 1.5-fold over AF568-TCO. These results identified T1 as the strongest neuron-associating ligand candidate in the screen and supported its use with P55 in dual-display PERCEPT formulations.

### Covalent PERCEPT display improves functional editing in neural progenitor cells

We next investigated whether covalent display of delivery and targeting ligands on the RNP surface improved functional genome editing as compared to the previously-reported “cocktail” format of PERC. Neural progenitor cells (NPCs) derived from the Ai9 mouse (which bears a CRISPR-compatible tdTomato fluorescent reporter cassette^35^) were treated with Cas9 RNP formulations and analyzed by flow cytometry.

To distinguish simple peptide admixture from covalent surface display, each ligand set was evaluated in three formats: admixed with unmodified RNP, admixed with linker-bearing RNP after the TCO moiety was inactivated via capping (“tz-cap”), or in the PERCEPT configuration: covalently conjugated to linker-decorated RNP. Unconjugated RNP and linker-only RNP produced low reporter activation, with ∼3–4% tdTomato-positive cells. TET1 alone provided limited benefit when covalently displayed, producing 5.7% tdTomato-positive cells. In contrast, admixed P55 increased editing to 14.5%, and covalent P55 tethered to RNP, or PERCEPT_P55_ further increased editing to 23.1%. Combining P55 with T1 produced the strongest gain, with covalent PERCEPT_P55,T1_ reaching ∼30.1% tdTomato-positive cells. Addition of 5 kDa PEG did not further increase editing, but qualitatively improved formulation handling and solubility (data not shown). Capping linker-bearing RNPs before ligand addition reduced editing toward admixed or lower-activity conditions, despite carrying matched peptide concentrations, providing evidence for the conjugation aspect enhancing functional delivery. Fluorescence microscopy of reporter activation corroborated the flow cytometry results (Fig. 1i,k). The T1-based enhancement of editing observed in NPCs did not occur in Ai9-derived fibroblasts (Extended Data Fig. 3), a finding consistent with our intended neuron-specific targeting.

To examine the entry mechanism of the optimized PERCEPT-RNP formulation, we treated Ai9 NPCs in the presence of endocytic inhibitors. Across 1×, 0.5× and 0.25× of the maximal dose, dynamin blockade with Dynasore, clathrin inhibition with Pitstop 2, and macropinocytosis inhibition with EIPA·HCl also yielded tdTomato-positive fractions comparable to DMSO controls, suggesting that none of these pathways were essential for PERCEPT activity^36^. In contrast, cholesterol sequestration with Filipin III dramatically attenuated editing at 1× and 0.5×, with partial recovery at 0.25×. Non-treated controls remained at background. These data show filipin-sensitive editing, consistent with a cholesterol-dependent caveolin mediated endocytosis delivery process for PERCEPT-RNP in NPCs (Extended Data Fig. 2).

### PERCEPT modulates tissue distribution and neuronal editing after striatal CED

We next asked whether PERCEPT surface programming could tune RNP delivery *in vivo*, where extracellular transport, cellular uptake, and cell-type engagement impose distinct barriers. Cas9 RNP formulations were administered by striatal convection enhanced delivery (CED)^37^ into Ai9 mice, and reporter activation was quantified across serial coronal sections. Unconjugated and linker-only RNP produced minimal tdTomato activation that was largely restricted to the needle track (Fig. 2b,c).

**Figure 2.**
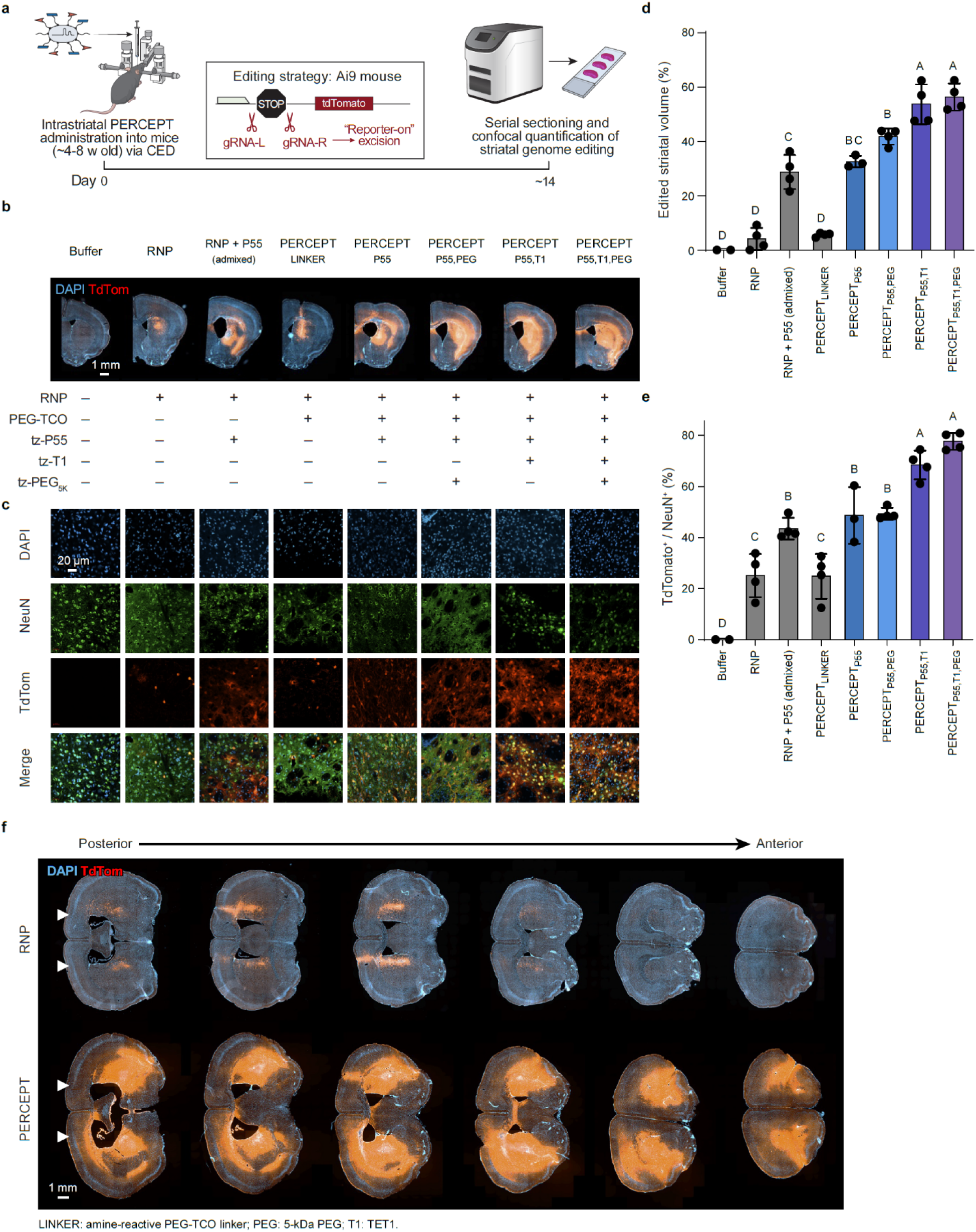
PERCEPT surface programming enhances local genome editing after striatal CED. a, Experimental design for striatal editing in Ai9 reporter mice. Cas9 RNP formulations were administered by intrastriatal convection-enhanced delivery, and genome editing was assessed by tdTomato reporter activation across serial brain sections. Brains were collected 14 days after administration and analyzed by cryosectioning and histology. **b,** Representative coronal sections showing tdTomato activation across the striatum after delivery of the indicated RNP formulations. **c,** Representative high-magnification immunofluorescence images from edited regions showing DAPI, NeuN and tdTomato signal. **d,** Quantification of edited striatal volume, calculated as tdTomato-positive volume divided by total striatal mask volume. **e,** Quantification of neuronal editing within the edited region, measured as the fraction of NeuN-positive cells that were tdTomato-positive. **f,** Representative serial coronal sections from control and optimized PERCEPT-treated brains, showing broader anterior-posterior distribution of tdTomato activation after optimized surface programming. PERC denotes RNP admixed with P55. Points represent hemispheres (four hemispheres from two mice per group). Letters show compact letter displays from ANOVA with Tukey’s test; groups sharing a letter are not significantly different. Scale bars, 1 mm (b,f) and 20 μm (c).

Admixing RNP with the amphiphilic delivery peptide P55 broadened reporter activation, editing approximately 28% of the striatal volume. Covalent attachment of P55 produced a modest additional gain. Addition of 5 kDa PEG increased edited striatal volume to approximately 41% without substantially changing neuronal editing efficiency, indicating that PEG primarily improved tissue editing distribution rather than productive engagement with neurons. Incorporation of the neuron-targeting ligand TET1 further increased edited volume to approximately 53% and increased neuronal editing, with approximately 68% of NeuN^+^ cells within the edited region becoming tdTomato-positive. The combined PERCEPT_P55,TET1,PEG_ formulation produced the broadest and most neuronally active editing profile, editing approximately 56% of the striatum and labeling approximately 78% of NeuN^+^ cells within the edited region (Fig. 2d,e). Multiplication of these two values suggests editing of ≥40% of all striatal neurons via PERCEPT_P55,TET1,PEG_ (Extended Data Fig. 11).

Whole-brain fluorescence imaging confirmed that optimized PERCEPT formulations produced broader reporter activation radiating from the injection site than control formulations. Representative serial coronal sections showed contiguous edited volumes across the striatum, and high-magnification images demonstrated robust tdTomato activation in NeuN^+^ neurons (Fig. 2b,c,f). Quantification was performed across serial sections from four hemispheres per group, corresponding to two mice per condition. Edited striatal volume was calculated as tdTomato-positive volume divided by total striatal mask volume. Together, these data indicate that PEG, P55 and TET1 contribute separable functions to local CNS delivery: PEG enhances tissue distribution, P55 promotes functional uptake, and TET1 increases neuronal engagement within the edited field.

### PERCEPT supports local neuronal editing in a distinct CNS region, the thalamus

To test whether PERCEPT-enhanced CNS delivery could be used in other regions of the brain, we applied the lead CNS formulation by CED into the thalamus of Ai9 mice. Unconjugated RNP produced only sparse tdTomato activation near the infusion site (Fig. 3a–d). In contrast, PERCEPT-formulated RNP generated broader reporter activation extending across multiple anterior-posterior thalamic sections (Fig. 3a–d; Extended Data Fig. 4). High-magnification imaging showed tdTomato signal within NeuN^+^ regions (Fig. 3b), consistent with neuronal editing in thalamic parenchyma. Quantification across serial sections showed increased edited thalamic tissue volume (Fig. 3c) and increased NeuN^+^ neuronal editing after PERCEPT delivery (Fig. 3d). These data indicate that PERCEPT-enhanced local CNS editing is not restricted to the striatum and can support neuronal editing in a second compact brain region.

**Figure 3.**
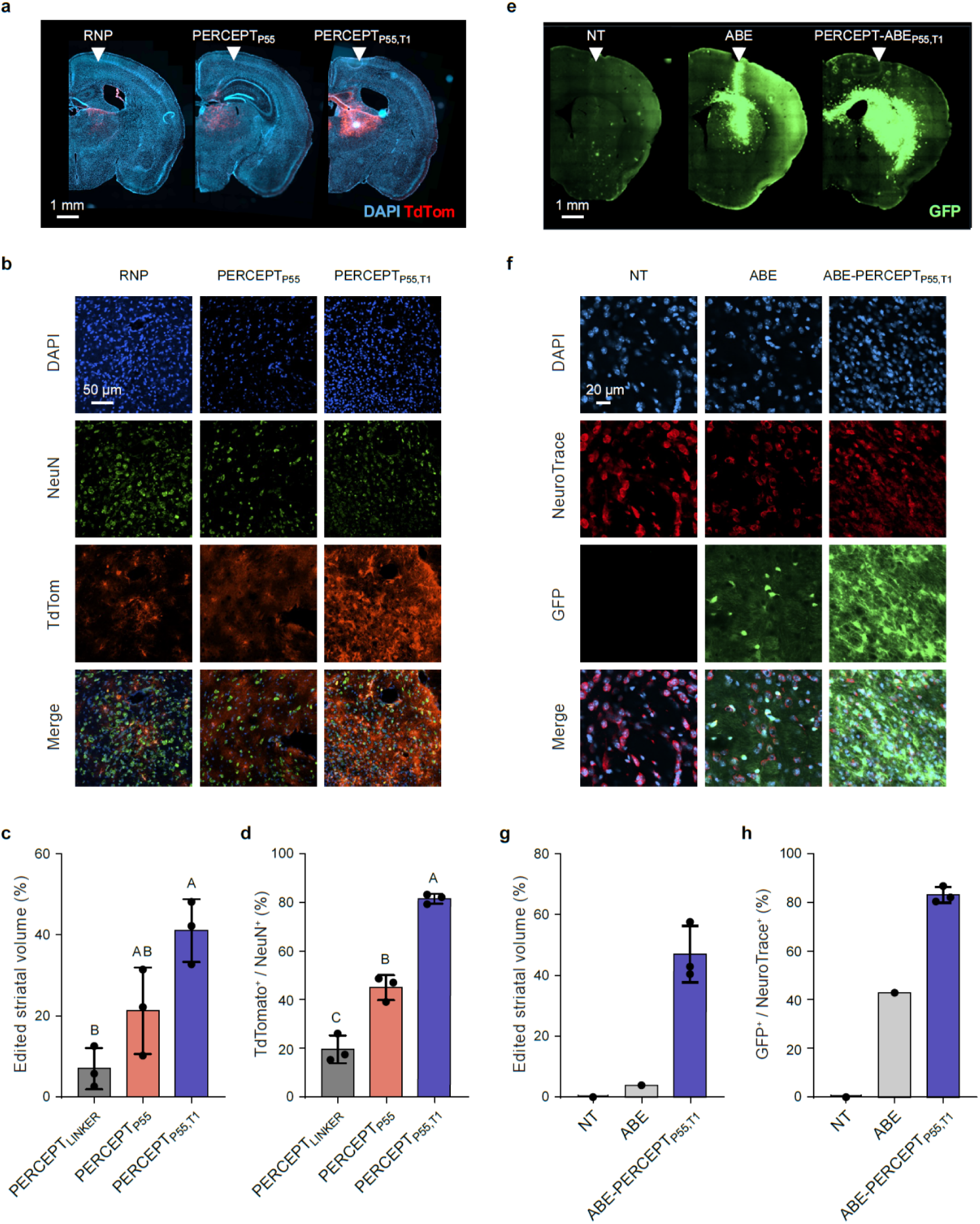
PERCEPT surface programming enhances local CNS genome editing in a second brain sub-structure and extends to adenine base editor delivery. a, Representative coronal brain sections from Ai9 tdTomato-on reporter mice after thalamic convection-enhanced delivery of Cas9 RNP formulations. White arrowheads indicate the approximate injection sites. PERCEPT-formulated RNP produced broader tdTomato reporter activation than control RNP formulations. **b,** Representative high-magnification immunofluorescence images from edited regions showing DAPI, NeuN, tdTomato and merged channels. **c,d,** Quantification of edited thalamic volume (c) and neuronal editing within the edited region (d). Neuronal editing was quantified as the fraction of NeuN-positive cells that were tdTomato-positive within the edited region. **e,** Representative coronal sections from GER10 reporter mice after striatal CED, showing GFP activation in untreated, unconjugated ABE and PERCEPT-ABE groups. White arrowheads indicate the approximate injection sites. **f,** High-magnification immunofluorescence images showing DAPI, NeuroTrace, GFP and merged channels. **g,h,** Quantification of GFP-positive striatal volume (g) and the percentage of NeuroTrace-positive cells that were GFP-positive (h). Reporter brains were collected 14 days after administration. Points represent hemispheres. For c,d, letters show compact letter displays from ANOVA with Tukey’s test; groups sharing a letter are not significantly different. NT, untreated. Scale bars, 1 mm (a,e), 50 μm (b) and 20 μm (f).

### PERCEPT supports adenine base editor RNP delivery

We next asked whether PERCEPT was compatible with other genome-editing cargo. An adenine base editor RNP^38^ was formulated for delivery into GER10 reporter mice^39^, wherein productive base editing activates GFP expression. At a matched dose and injection volume, convection enhanced delivery of unconjugated ABE RNP produced limited GFP activation centered near the infusion track (Fig. 3e). In contrast, PERCEPT-formulated ABE generated broader GFP reporter activation across the dorsal striatum, with signal extending beyond the injection site and across serial coronal sections (Fig. 3e; Extended Data Fig. 10). High-magnification imaging showed GFP-positive cells within NeuroTrace-rich regions, consistent with neuronal base editing after local delivery (Fig. 3f). Quantification across serial sections showed increased GFP-positive edited volume with PERCEPT-ABE relative to unconjugated ABE (Fig. 3g), as well as increased neuronal reporter activation (Fig. 3h). These findings suggest that PERCEPT can improve local CNS delivery of adenine base editor RNPs, extending the platform beyond conventional nuclease cargo.

### PERCEPT-RNP enables disease-relevant editing in a murine Huntington’s disease model

Having established PERCEPT-mediated CNS reporter editing, we next tested whether the lead CNS formulation could support disease-relevant editing in the R6/2 Huntington’s disease model^40^ (Fig. 4). Bilateral intrastriatal CED of PERCEPT_P55,TET1,PEG_ targeting the human HTT transgene produced guide-dependent editing in striatal tissue, whereas a GFP-guide control showed no detectable modification at the HTT target site (Fig. 4b). Targeted amplicon sequencing demonstrated ∼10% modified reads in bulk cells from HTT-targeted striata, with near-baseline levels in mock-treated striatum and cerebellar controls from the same animals. We note that the glia:neuron ratio in the murine striatum has been estimated as 3:1, so if PERCEPT exhibited effective neuronal tropism this editing efficiency would correspond to editing of ∼40% of all striatal neurons, consistent with the editing efficiency we observed in Ai9 reporter mice (Fig. 2d,e; Extended Data Fig. 11).

**Figure 4.**
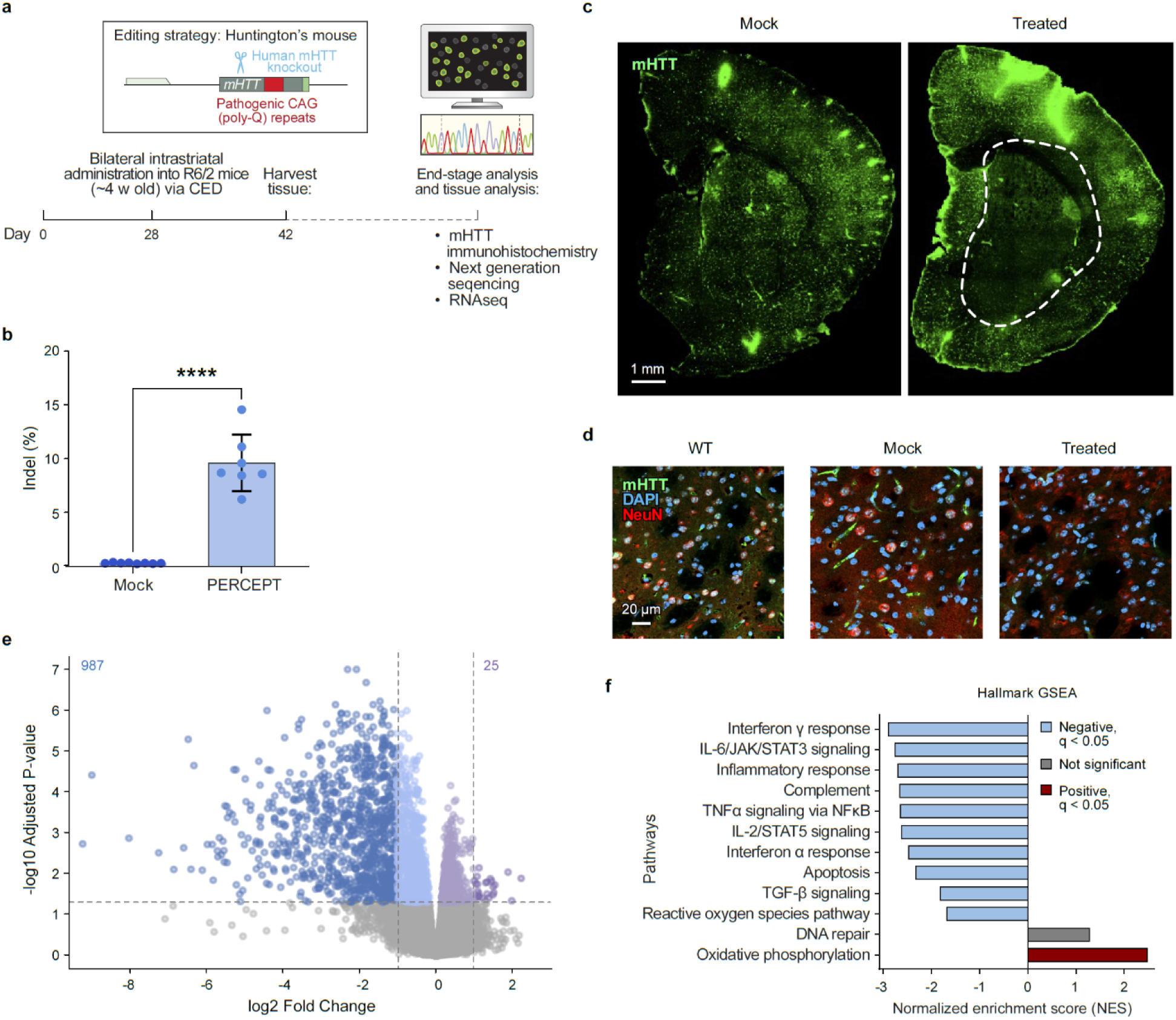
PERCEPT-mediated editing of mutant huntingtin in the R6/2 Huntington’s disease model reduces mHTT pathology and disease-associated transcriptional programs. a, R6/2 mice received bilateral intrastriatal convection-enhanced delivery of PERCEPT-formulated Cas9 RNP targeting the human mutant HTT transgene, or control GFP-targeted PERCEPT-formulated Cas9 RNP, followed by histological and molecular analysis. The schematic shows the editing strategy targeting the pathogenic human mHTT transgene containing expanded CAG/polyQ repeats. **b,** Targeted amplicon sequencing of dissected tissue showed approximately 10% indels in PERCEPT-treated samples and near-background indels in nonsense guide PERCEPT treated controls. **c,** Representative mHTT immunofluorescence images show reduced mutant huntingtin aggregate burden within the edited striatal region, outlined by a white dashed line, compared with surrounding or control tissue. **d,** High-magnification immunostaining shows mHTT, DAPI and NeuN signal in striatal sections from wild-type (WT), mock-treated and treated mice. **e,** Volcano plot of differential gene expression in treated versus sick control treated striata. Blue, significantly decreased; purple, significantly increased; grey, not significant. **f,** Hallmark gene set enrichment analysis. Bars show normalized enrichment scores; blue and red indicate significant negative and positive enrichment (q < 0.05), respectively. Grey indicates non-significant enrichment (DNA repair, q = 0.983). Scale bars, 1 mm (c) and 20 μm (d).

Immunostaining for mutant huntingtin revealed a qualitative reduction in aggregate burden within the edited striatal region, consistent with local disruption of the mutant HTT transgene (Fig. 4c,d). Transcriptomic profiling of treated striata showed suppression of HD-associated reactive glial and injury programs, including complement and inflammatory signatures elevated in control-treated tissue, consistent with reported glial alterations^41^, complement-associated pathology^42^ and neuroinflammation^43^ known to occur in HD. Gene set enrichment analysis (GSEA)^44^ showed reduced IL6-JAK-STAT3, TNFα/NFκB, interferon-response and complement pathway activity, alongside relative enrichment of oxidative phosphorylation and modest, non-significant DNA repair signatures in HTT-targeted samples. Together, these data show that PERCEPT-RNP delivery can produce localized, guide-dependent editing of disease-relevant mutant HTT *in vivo*, with accompanying reductions in mHTT aggregate burden and transcriptional changes consistent with reduced disease-associated inflammatory and injury signaling. Together these data show that PERCEPT-mediated RNP editing is accompanied by coordinated histological and transcriptional changes.

### PERCEPT edits retinal cells following intravitreal delivery

We next evaluated whether PERCEPT surface programming could be adapted to a second tissue with distinct anatomical barriers. Ai9 tdTomato-on reporter mice received intravitreal delivery of RNP formulations, and retinal sections were stained for Sox9 to quantify Müller glia editing^45^. Buffer and unconjugated RNP produced little tdTomato activation in Sox9-positive Müller glia. PERCEPT_P55_ increased reporter activation modestly. In contrast, dual conjugation with P55 and TET1 generated widespread tdTomato signal across the inner retina, with co-localization in Sox9-positive somata and radial processes characteristic of Müller glia (Fig. 5b,c). PERCEPT_P55,TET1_ produced approximately 53% tdTomato-positive Sox9-positive cells, compared with approximately 10% after PERCEPT_P55_ and near-baseline signal after buffer treatment.

**Figure 5.**
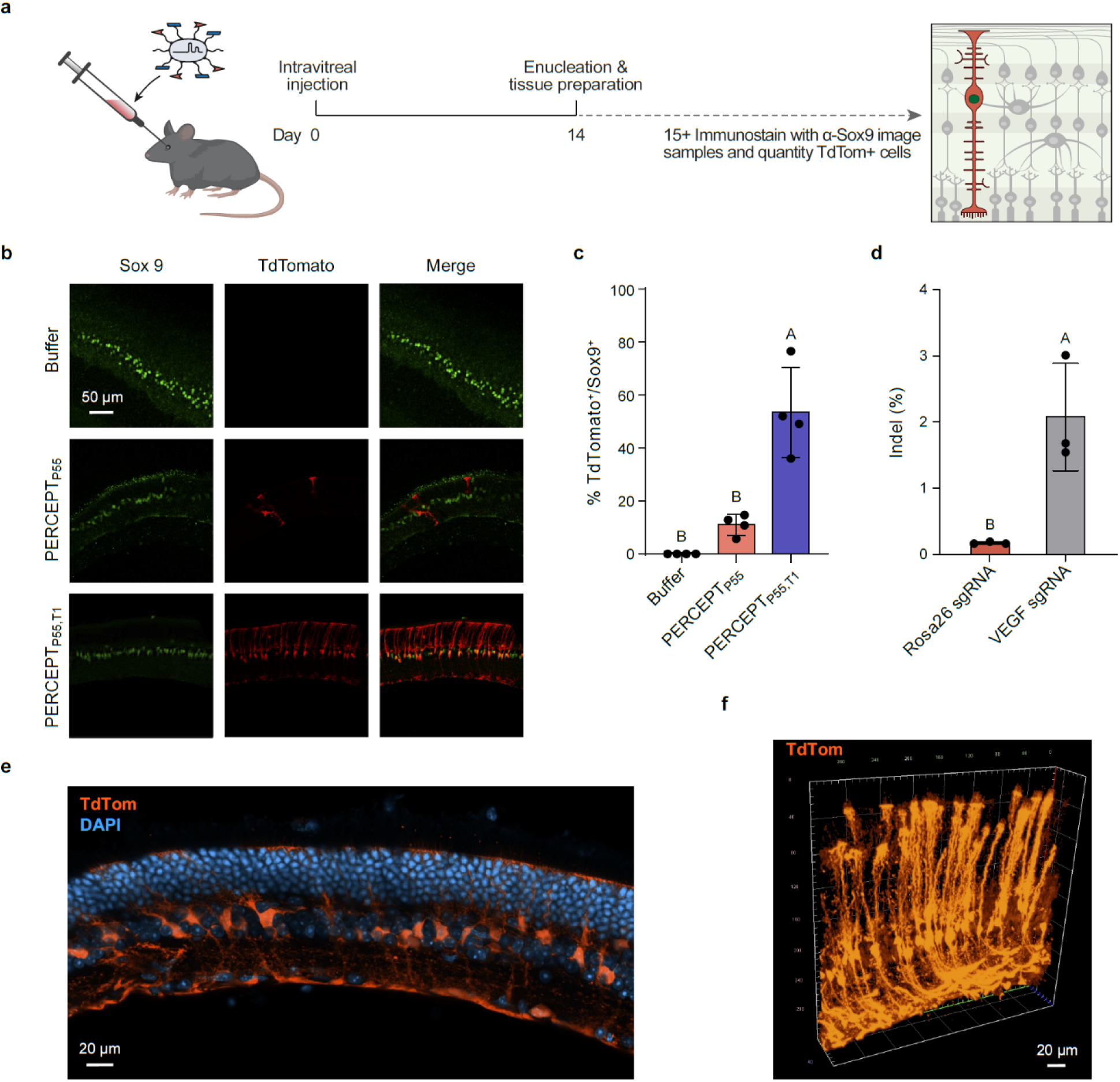
PERCEPT surface programming enables Müller glia editing after intravitreal delivery. a, Experimental design for intravitreal delivery in Ai9 tdTomato reporter mice. RNP formulations were administered by intravitreal injection on day 0. Eyes were enucleated and processed on day 14, then retinal sections were immunostained for Sox9 and imaged to quantify Müller glia editing. **b,** Representative retinal sections showing Sox9 immunostaining and tdTomato reporter activation after buffer, P55-containing PERCEPT or P55/TET1-containing PERCEPT treatment. Dual surface programming with P55 and TET1 generated broad tdTomato labeling across the inner retina, with signal extending through Sox9-positive Müller glial somata and radial processes. **c,** Quantification of tdTomato-positive Sox9-positive cells after buffer, P55-containing PERCEPT or P55/TET1-containing PERCEPT treatment. **d,** Targeted amplicon sequencing of whole-retina lysates after intravitreal delivery of the optimized PERCEPT formulation with a VEGF-targeting guide or a Rosa26 control guide. **e,** Representative confocal image of retinal tissue showing tdTomato-positive somata and radial processes. **f,** Three-dimensional reconstruction of a confocal z-stack showing tdTomato-positive radial processes. Points represent individual eyes. Letters show compact letter displays from ANOVA with Tukey’s test; groups sharing a letter are not significantly different. Scale bars, 50 μm (b) and 20 μm (e,f).

To test whether this delivery activity extended to an endogenous locus, we performed a parallel intravitreal study using a VEGF-targeting guide. Targeted amplicon sequencing detected approximately 1.5% edited reads across whole-retina lysates, while control-treated samples remained near baseline. Because sequencing was performed from whole-retina lysates, the bulk editing value likely underestimates editing within the responsive Sox9-positive Müller glia population, which only represent ∼3% of cells in the murine retina^46^. Together, these data show that dual P55/T1 surface programming enables efficient Müller glia reporter editing and measurable endogenous-locus editing after intravitreal RNP delivery.

### Optimal skeletal muscle editing requires a distinct PERCEPT formulation

We next asked whether PERCEPT surface chemistry could be adapted for local delivery in skeletal muscle, a dense peripheral tissue composed of large multinucleated fibers. Following bilateral intramuscular injection of 50 µL of 40 µM RNP per gastrocnemius, unconjugated RNP produced little tdTomato activation, while PERCEPT_P55_ yielded labeling that remained largely proximal to the needle track. Substituting P55 with the alternative amphiphilic peptide S315^47^ increased gastrocnemius fluorescence by approximately fivefold relative to PERCEPT_P55_, indicating that peptide identity strongly influenced functional delivery in muscle. Addition of 5 kDa PEG further improved tissue distribution: PERCEPT_P55,PEG_ broadened tdTomato signal distal to the needle track and increased fluorescence relative to PERCEPT_P55_, while PERCEPT_S315,PEG_ produced the strongest overall response, reaching approximately tenfold higher fluorescence than unconjugated RNP and generating broad fields of tdTomato⁺ myofibers spanning large fascicles (Fig. 6b,d,f). Whole-organ epifluorescence localized reporter signal to the injected gastrocnemius, and section imaging confirmed tdTomato activation in myofiber-like structures (Extended Data Fig. 5). Targeted amplicon sequencing also detected editing at the endogenous myostatin locus in gastrocnemius muscles treated with PERCEPT containing S315 and PEG (Fig. 6e). Thus optimal muscle delivery requires a PERCEPT surface chemistry distinct from the lead CNS formulation, with S315 and PEG combining to improve local myofiber editing and tissue spread.

**Figure 6.**
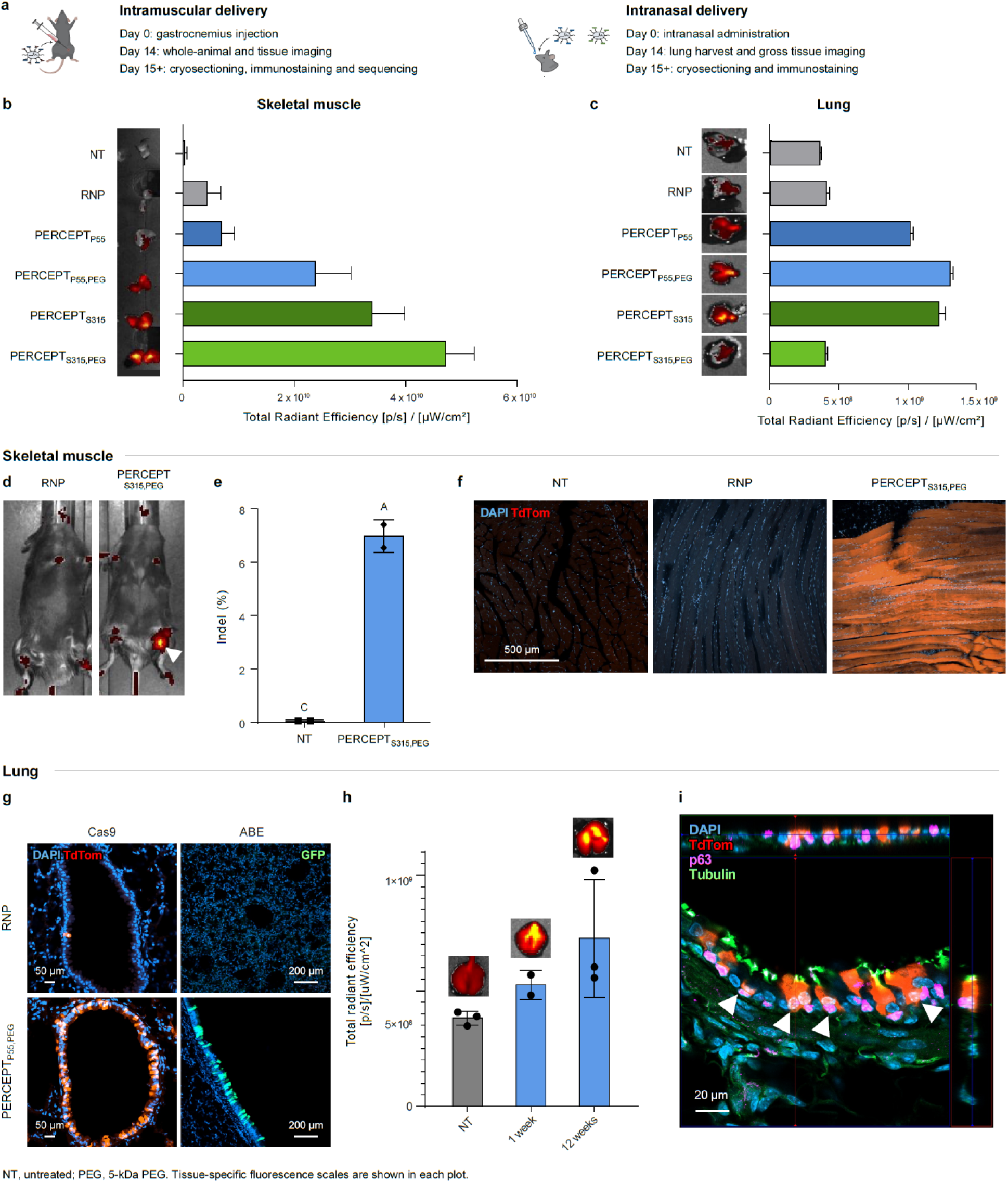
PERCEPT surface programming enables local genome editing in skeletal muscle and lung. a, Intramuscular/intranasal dosing, day 0; day 14, whole-animal fluorescence imaging (muscle study) and gastrocnemius/lung harvest for gross imaging; from day 15, cryosectioning/immunostaining and muscle sequencing. Lung durability timepoints, h. **b,c,** *Ex vivo* gastrocnemius (b) and whole-lung (c) fluorescence images and total radiant efficiency ([p/s]/[μW/cm²]) for indicated formulations: three separate image acquisitions (b); repeated acquisitions of the same lung (c). PERCEPT_P55,PEG_ produced the strongest whole-lung signal. **d,** Whole-animal fluorescence imaging after unconjugated RNP or PERCEPT_S315,PEG_. **e,** Myostatin indel frequencies in NT and PERCEPT_S315,PEG_: Quintara Illumina sequencing, two gastrocnemius muscles/group. PERCEPT used a myostatin-targeting guide (Extended Data Table 2). **f,** Representative gastrocnemius cryosections showing DAPI/tdTomato: untreated, unconjugated RNP and PERCEPT_S315,PEG_. Optimized PERCEPT generated broad tdTomato-positive myofiber labeling across the injected region. **g,** Representative lung sections showing tdTomato activation after intranasal Cas9 RNP (Ai9 mice) or GFP activation after ABE (GER10 mice), concentrated around conducting airways. Upper/lower rows: unconjugated RNP/PERCEPT_P55,PEG_. **h,** Representative lung fluorescence images and total radiant efficiency at 1 and 12 weeks after dosing, with untreated controls; each point represents a different animal. Signal remained detectable at 12 weeks. **i,** Orthogonal confocal views (DAPI, tdTomato, p63, tubulin) support tdTomato localization within p63-positive basal cells adjacent to tubulin-positive cells. For e, compact letter displays from ANOVA with Tukey’s test; shared letters indicate no significant difference. NT, untreated; PEG, 5-kDa PEG. Scale bars, 500 μm (f), 50 μm (g, Cas9), 200 μm (g, ABE), 20 μm (i).

### PERCEPT mediates durable editing of the airway epithelium

Finally, we tested whether PERCEPT surface programming could support local RNP delivery to the lung after intranasal administration. Ai9 mice received 25 µL of 20 µM RNP formulation via intranasal administration. Although unconjugated and linker-only RNP produced little tdTomato activation in the lung, PERCEPT_P55_ generated patchy reporter signal adjacent to conducting airways, whereas PEGylation substantially improved pulmonary editing. PERCEPT_P55,PEG_ produced strong, continuous peribronchial tdTomato signal consistent with airway epithelial labeling and emerged as the lead pulmonary formulation. PERCEPT_S315_ and PERCEPT_S315,PEG_ also generated reporter activation, but PERCEPT_P55,PEG_ produced the strongest overall airway-associated signal and was prioritized for follow-up studies. Whole-animal and excised-organ epifluorescence localized reporter activation primarily to the respiratory tract, with minimal detectable signal in non-respiratory tissues (Extended Data Fig. 6). Quantification of whole-lung fluorescence confirmed increased reporter signal after optimized PERCEPT delivery relative to minimally active control formulations (Fig. 6c). Whole lung tdTomato signal remained detectable and stable up to 12 weeks after dosing (Fig. 6h), demonstrating durable genomic editing after local intranasal RNP delivery.

Histological analysis showed strong tdTomato labeling concentrated around conducting airways. Reporter activation was observed in airway epithelia, including p63-positive basal cells and adjacent ciliated cells. Orthogonal confocal views supported localization of tdTomato within p63-positive cells, suggesting Basal cell progenitor editing^48^ (Fig. 6i; Extended Data Fig. 7). An analogous adenine base editor formulation produced GFP reporter activation in airway-associated regions, indicating that PERCEPT surface programming can also support pulmonary delivery of larger editor RNP cargos (Fig. 6g). Together, these data identify PERCEPT_P55,PEG_ as a lead PERCEPT formulation for durable pulmonary genome editing via intranasal administration.

## Discussion

By combining a redox-sensitive self-immolative linker with PEG-mediated shielding and a bioorthogonal click handle for rapid ligand decoration, PERCEPT converts Cas9 and adenine base editor RNPs into compact, tunable delivery scaffolds without permanently altering the genome-editing payload. Material characterization confirmed release under reducing conditions, while conjugated particles remained small relative to other nonviral carriers like LNPs, and conceptually more compatible with transport through extracellular spaces.

A central finding of this study is that individual PERCEPT surface components contribute separable delivery functions *in vivo*. In the striatum, PEGylation broadened the edited region without substantially increasing the fraction of edited neurons within that region, consistent with improved local distribution^11,24^. By contrast, P55 and TET1 increased functional cellular engagement, with TET1 increasing neuronal editing within the edited field. The combined P55/TET1/PEG formulation therefore joined broader tissue coverage with stronger neuronal editing. In vitro, covalent ligand display increased activity relative to unconjugated or admixed controls, whereas PEG improved formulation handling without increasing editing. Together, these data suggest that PEG improves spatial access, amphiphilic peptides promote productive intracellular delivery, and targeting ligands bias editing toward selected cell types.

This surface-programming logic was not confined to one anatomical site or editor architecture. The lead CNS formulation supported local neuronal editing in the thalamus and improved delivery of an adenine base editor RNP in GER10 reporter mice, demonstrating that the delivery scaffold can be separated from the editing payload. In the R6/2 Huntington’s disease model, PERCEPT-formulated RNPs enabled guide-dependent editing of the human mutant HTT transgene, reduced mutant huntingtin aggregate burden within the edited region, and suppressed reactive glial, inflammatory, complement, and injury-associated transcriptional programs. These findings extend PERCEPT beyond reporter activation and show that local RNP delivery can produce disease-relevant editing accompanied by histological and transcriptional changes consistent with reduced HD-associated inflammatory and injury signaling.

The same modular chemistry generalized beyond the CNS, but the preferred surface chemistry changed with tissue context. Dual P55/TET1 display enabled Müller glia reporter editing and detectable endogenous VEGF editing after intravitreal delivery. In skeletal muscle, use of S315 and PEG moieties produced broader and stronger myofiber editing than the lead CNS delivery peptide P55. In the lung, a P55+PEG formulation emerged as the lead formulation after intranasal administration, producing airway-associated reporter activation that remained detectable for 12 weeks. Co-staining supported editing within airway epithelial populations, including p63-positive basal cells, and a parallel adenine base editor formulation produced airway-associated reporter activation. These tissue comparisons suggest that PERCEPT provides a screening-friendly framework for matching RNP surface chemistry to local biological environments.

Future work will quantify editing at disease-relevant loci in the target-cell populations and determine how surface modifications affect cellular uptake and endosomal escape. The present study does not thoroughly characterize *in vivo* tolerability or characterize off-target editing. Further studies will assess these outcomes, together with editing durability and the effects of repeated or higher-dose administration, as PERCEPT is developed for clinically relevant delivery routes and therapeutic applications.

Taken together, PERCEPT establishes a cleavable, click-enabled surface-programming strategy that can tune where an RNP editor goes *in vivo* while preserving a compact and transient delivery scaffold. The highest-performing chemistry may depend on local anatomical and cellular context, but we were encouraged to find that the same framework was cross-compatible nuclease and base editor cargos – suggesting plug & play use with other genome editing enzymes – and supported disease-relevant editing in the R6/2 model.

## Methods

### Animals and study design

Animal experiments were approved by the University of California, Berkeley Animal Care and Use Committee under Ross Wilson’s protocol AUP-2018-08-11339-2. Ai9 tdTomato reporter mice were used for Cas9 reporter-editing studies,^35^ GER10 Venus/GFP reporter mice for CNS and pulmonary adenine base editor (ABE) studies,^39^ and R6/2 mice for human HTT transgene editing.^40^ Published stock identifiers are 007909 for Ai9 and 069724-JAX for GER10.^49^ Animals were randomly allocated to treatment groups, and investigators were blinded where feasible. In reporter studies, fluorescence frequently revealed treatment assignment after tissue collection, limiting blinding during subsequent assessment. R6/2 treatment assignments remained blinded until sequencing. Mice were group-housed on a 12-h light–dark cycle at 21 ± 2°C and 50–60% humidity, with food and water available ad libitum, similar to our previous work.^49^

For stereotactic, intramuscular and intranasal administration, mice were maintained under 2% isoflurane anesthesia with thermal support at 37°C and ocular lubrication.^18,49,50^

### Peptides and linker synthesis

Peptides were synthesized commercially by CPC Scientific using fluorenylmethyloxycarbonyl (Fmoc) solid-phase chemistry on Rink amide or Wang resin and cleaved using a trifluoroacetic acid cocktail. Peptides were purified by reversed-phase high-performance liquid chromatography (HPLC); purity and identity were assessed by analytical HPLC and electrospray mass spectrometry, respectively. Peptide sequences and conjugation handles are listed in Extended Data Table 1.

The PERCEPT linker, PNC–DS–PEG23–TCO (TCO–PEG23–DEC-lite), comprised an activated para-nitrophenyl carbonate, a reduction-sensitive disulfide, a PEG spacer containing 23 ethylene glycol units and a trans-cyclooctene (TCO) handle. Amine-terminated TCO–PEG23 (0.34 mmol) and triethylamine (523 µL, 3.77 mmol) in dimethylformamide (5 mL) were added dropwise over 15 min at 0°C to the activated disulfide carbonate (1.65 g, 3.4 mmol) in dimethylformamide (10 mL). The reaction was warmed to room temperature and stirred overnight. After solvent removal, the residue was dissolved in dichloromethane (50 mL), washed with 0.5 M HCl (2 × 30 mL) and brine (30 mL), and dried over sodium sulfate. Preparative thin-layer chromatography (dichloromethane/methanol, 10:1) yielded 305 mg (57%) of linker as a waxy white solid. Proton NMR spectra were acquired from 2 mg linker in 600 µL CDCl_3_ using a Bruker AVANCE 400 spectrometer with an Oxford Instruments 9.4 T magnet at the UC Berkeley College of Chemistry NMR Facility (Extended Data Fig. 9). The separately conjugated shielding polymer was methoxy-PEG–methyltetrazine of nominal molecular mass 5 kDa (BroadPharm, BP-26353).

The activated disulfide carbonate was prepared by adding p-nitrophenyl chloroformate (30 g, 148.8 mmol) portionwise to bis(2-hydroxyethyl) disulfide (7.6 g, 49.6 mmol) and pyridine (15.6 mL, 198.4 mmol) in dichloromethane (150 mL) in an ice bath. The mixture was warmed to room temperature and stirred overnight, diluted with dichloromethane (100 mL), and washed with 0.5 M HCl (3 × 50 mL), sodium bicarbonate (5 × 50 mL) and brine (3 × 50 mL). Drying over sodium sulfate, concentration and recrystallization from diethyl ether yielded 16.5 g (68.5%) of white powder.

To prepare the amine-terminated TCO–PEG23 precursor, TCO NHS carbonate (137 mg, 0.512 mmol), Boc-protected diamino-PEG23 (500 mg, 0.426 mmol) and triethylamine (118 µL, 1.278 mmol) were stirred in dry dimethylformamide (10 mL) under nitrogen at room temperature overnight. After evaporation, the residue was dissolved in dichloromethane (50 mL), washed with 0.5 M HCl (2 × 10 mL), sodium bicarbonate (2 × 10 mL) and brine (10 mL), and dried over sodium sulfate. Silica chromatography (Biotage Selekt; dichloromethane/methanol, 0–10%) yielded 450 mg (80%) of Boc-protected TCO–PEG23 as a colorless oil. This intermediate (450 mg, 0.343 mmol) was treated with trifluoroacetic acid (4 mL) in dichloromethane (20 mL) in an ice bath for 3 h. Toluene (5 mL) was added before solvent removal; the residue was redissolved in dichloromethane/toluene (10/5 mL) and concentrated again to remove residual acid.

### CRISPR proteins and guide RNAs

Recombinant *Streptococcus pyogenes* Cas9 carried three nuclear localization signals (tri-NLS) and the C80S/C574S substitutions (Addgene, 196244). The ABE8e editor was assembled from SpyTag-tagged TadA8e and SpyCatcher–nCas9, as described previously.^49^ Proteins were expressed in *Escherichia coli* and purified by nickel-affinity, ion-exchange and size-exclusion chromatography. TadA8e and nCas9 were expressed in BL21 STAR(DE3) and Rosetta(DE3), respectively. Purified components were combined for 30 min at room temperature, followed by Superdex S200 size-exclusion purification. Proteins were concentrated to approximately 50 µM and stored at −80°C in 20 mM HEPES–KOH, 150 mM NaCl and 10% glycerol, pH 7.5.

Synthetic single-guide RNAs were obtained from Synthego or IDT, dissolved in water and refolded in 20 mM HEPES, pH 7.5, and 150 mM NaCl by heating at 95°C for 5 min and cooling to room temperature, before adding to Cas9 protein at a 1.5:1 (Guide:Cas9) ratio. Guide spacer sequences and vendors are provided in Extended Data Table 2.

### RNP assembly and surface functionalization

Editor proteins and guide RNAs were combined in RNP buffer containing 20 mM HEPES, pH 7.4, 150 mM NaCl, 200 mM trehalose and 2 mM MgCl_2_. DMSO-solubilized DEC linker (PNC-DS-PEG-TCO) was added after RNP assembly at an 80:1 linker:RNP molar ratio. Samples were mixed at approximately 300 rpm for 4 h at 22°C, cooled to 4°C and washed three times in 50 kDa molecular weight cutoff (MWCO) Amicon centrifugal filter devices.

Tetrazine-bearing peptides were mixed with dilution buffer and linker-modified RNP and reacted for 1 h at 22°C. Samples were used immediately or flash-frozen at −80°C and thawed at room temperature.

Formulations included P55, TET1/T1, S315 and 5-kDa PEG, singly or in combinations used in each experiment. Comparators included unmodified RNP, linker-only RNP, peptide-admixed RNP, and linker-bearing RNP capped before peptide addition.

### Reductive cleavage analysis

Unmodified, linker-modified and peptide-conjugated SpCas9 RNPs were compared by denaturing SDS–PAGE under reducing and non-reducing conditions. Reducing samples contained 2.5% β-mercaptoethanol per sample and were heated at 98°C for 5 min. Samples were resolved on Bio-Rad 4–20% Tris gels at 250 V for 23 min.

### Particle size measurement

Hydrodynamic size was measured by dynamic light scattering using a Zetasizer Nano ZS (Malvern Panalytical) and a low-volume quartz cuvette (ZEN2112). Samples were mixed by pipetting, equilibrated for 1 min and measured at 25°C, with buffer blanks and three technical readings per sample. Data were processed using Zetasizer software, and size distribution reported by number.

### Electron microscopy

For negative-stain transmission electron microscopy, formvar-coated copper grids were dipped into 0.4 µM RNP preparations. Excess liquid was blotted, and grids were stained with 2% uranyl acetate and air-dried. Images were acquired using a Tecnai 12 transmission electron microscope (FEI) at accelerating voltages of 80–120 kV.

Scanning electron microscopy was performed on carbon-sputtered PERCEPT samples using a Crossbeam 550 (Zeiss) at 0.7 kV, with a working distance of 4.3 mm and an SESI detector.

### Cell culture

Ai9 neural progenitor cells (NPCs) were the previously described line derived from cortical tissue of homozygous Ai9 mouse embryos at embryonic day 13.5 using the papain-based Neural Dissociation Kit (Miltenyi Biotec, 130-092-628)^50^. Cells were maintained as non-adherent neurospheres in DMEM/F12 (Thermo Fisher Scientific, 10565-018), B-27 without vitamin A (12587-010), N-2 (17502-048), non-essential amino acids (11140-050), 10 mM HEPES, 2-mercaptoethanol (21985-023) and penicillin–streptomycin (15140-122). Basic FGF (BioLegend, 579606) and EGF (Thermo Fisher Scientific, PHG0311) were each supplied at 20 ng/mL. Cultures were maintained at 37°C and 5% CO_2_; growth factors were refreshed every 3 days.

Neurospheres were dissociated every 6 days and replated at approximately 1.5 million cells per 10-cm dish. Cells were used between passages 2 and 15. Cultures were tested weekly for mycoplasma using the MycoAlert Mycoplasma Detection Kit (Lonza); all tests were negative.

Neuro2A cells (ATCC, CCL-131) were cultured according to the ATCC protocol in Eagle’s minimum essential medium (ATCC, 30-2003) containing 10% fetal bovine serum at 37°C in 5% CO_2_.^51^

### Fluorescent peptide screening

Tetrazine-modified candidate peptides were coupled to AZDye 568 TCO (Click Chemistry Tools, 1358-5) for 1 h at room temperature. Free dye was not removed before the labelled peptide mixtures were applied to Neuro2A cells. Cell-associated fluorescence was measured in gated cell populations using an Attune flow cytometer (Thermo Fisher Scientific) with four fluorescence channels and normalized to dye alone. Selected treatment groups were imaged on a Zeiss LSM 900 confocal microscope using a 20× objective.

### NPC editing and viability

Ai9 NPCs were plated at 5,000 cells per well in 96-well plates coated with poly-DL-ornithine (Sigma-Aldrich, P8638), laminin (11243217001) and fibronectin (F4759), and were treated with Cas9 RNP formulations targeting the tdTomato reporter, similar to previous work^18,50^. After 72 h, cells were washed with PBS, detached using 0.25% trypsin, neutralized with DMEM containing 10% fetal bovine serum, and resuspended in PBS containing 1% fetal bovine serum and EDTA.

Flow cytometry used an Attune NxT (Thermo Fisher Scientific). Data were analyzed in Attune software, with untreated cells defining the negative control and editing measured as the percentage of tdTomato-positive cells. Cell viability was assessed using the Fixable Violet Dead Cell Stain Kit (Thermo Fisher Scientific, L34964; 1:1,000) for 20 min at 4°C in the dark.^49^ Viability was determined from the dye-negative live-cell gate and compared across formulations and vehicle controls.

Reporter activation was also assessed using a Revolve widefield fluorescence microscope (Echo), as described previously.^50^

### Endocytic inhibitor treatment

Ai9 NPCs received 40 pmol PERCEPT RNP for 90 min in the presence of Dynasore (Cell Signaling Technology, 46240S), Pitstop 2 (Sigma-Aldrich, SML1169)^36^, EIPA hydrochloride (Fisher Scientific, 50-196-7883) or Filipin III (Sigma-Aldrich, SAE0087), with DMSO and untreated controls. The 1× concentrations were 30 µM Dynasore, 60 µM Pitstop 2, 200 µM EIPA hydrochloride and 4 µg/mL Filipin III; the 0.5× and 0.25× conditions used one-half and one-quarter of these concentrations. Reporter activation was measured by flow cytometry.

### Striatal and thalamic delivery

Cas9 RNP formulations were administered by convection-enhanced delivery into the striatum or thalamus of Ai9 mice. ABE formulations were delivered into the dorsal striatum of GER10 mice, with dose and volume matched between unconjugated and PERCEPT groups. CNS formulations contained 40 µM RNP. Striatal delivery adapted the procedure described previously.^49^ Anaesthetized mice received subcutaneous buprenorphine and were positioned in a stereotactic frame. A stepped fused-silica cannula (1 mm step) targeted AP +0.8 mm and ML ±2.0 mm relative to bregma, at DV −2.6 mm below the skull surface. A microsyringe pump (World Precision Instruments) delivered 5 µL per hemisphere for striatum (200 pmol RNP) or 2 µL per hemisphere for thalamus (80pmol) at 0.5 µL/min; the cannula remained in place for 10 min before withdrawal. Mice were monitored through recovery and twice daily for 5 days. Ai9 striatal reporter tissues were collected 14 days after dosing.

### CNS histology and reporter quantification

Brains for histology were processed as described previously.^49^ Mice were transcardially perfused with PBS followed by 4% paraformaldehyde, pH 7.4. Brains were post-fixed for 24 h, cryoprotected in 30% sucrose in PBS until sectioning, embedded in OCT and stored at −80°C. Reporter cohorts used 20-µm serial coronal sections mounted on slides. Sections were washed in PBS containing 0.3% Triton X-100 and blocked in 1% bovine serum albumin and 5% normal goat serum. Neurons were labelled with chicken anti-NeuN (GeneTex, GTX00837) or NeuroTrace (Invitrogen N21483). Primary antibodies were applied overnight at 4°C and matched secondary antibodies for 1 h at room temperature, with PBS washes between steps. Sections were counterstained with DAPI (1:5,000; MilliporeSigma, MBD0015) and mounted in Fluoromount (MilliporeSigma, F4680).

For Ai9 studies, the tdTomato-positive tissue fraction was calculated as tdTomato-positive volume divided by total striatal or thalamic mask volume. Neuronal editing was calculated as the fraction of NeuN-positive cells that were tdTomato-positive within the edited region. Serial images were processed in MATLAB for alignment, segmentation and volume reconstruction.

GER10 whole sections were imaged on a Zeiss Axio Scan 7 using a 20× objective with standardized exposure and gain settings, and exported as 16-bit multichannel TIFF files. Confocal imaging used a Zeiss LSM 990 with a Plan-Apochromat 20× objective and sequential acquisition of DAPI, GFP and NeuroTrace far-red channels. NeuN immunostaining was used for neuronal classification in the quantitative analysis.

Tissue coverage was analyzed in MATLAB. Within each anatomical region of interest, tdTomato or GFP signal was segmented using Otsu thresholding. Morphological opening and removal of small objects, followed by hole filling and convex-hull processing, generated the reporter distribution map. Neuronal editing percentages were determined manually from five randomly selected confocal images acquired at 20× within the edited striatal region. The five image-level percentages were averaged to yield one data point per hemisphere.

### Huntington disease experiments

R6/2 mice (3–4 weeks old, approximately 160 CAG repeats) received bilateral intrastriatal PERCEPT containing P55, TET1 and 5-kDa PEG, carrying the human HTT-targeting SpHD1 guide (Extended Data Table 2) or a GFP-targeting control guide. SpHD1 was adapted for SpCas9 from the HTT-targeting strategy previously delivered using AAV and SaCas9.^49,52,53^ Delivery used the striatal dose, infusion rate and surgical procedure described above. Striatal tissue was analyzed by targeted amplicon sequencing, mutant huntingtin immunostaining and RNA sequencing. Cerebellar tissue and mock-treated striata were included as sequencing controls.

Mutant huntingtin immunostaining used 40-µm free-floating coronal sections with the fixation, blocking and antibody-incubation procedure described above.^49^ Primary antibodies were mouse anti-huntingtin mEM48 (MilliporeSigma, MAB5374; RRID:AB_177645; 1:100) and rabbit anti-NeuN (MilliporeSigma, ABN78; 1:100).

### Intravitreal delivery and retinal analysis

RNP formulations were administered intravitreally to Ai9 mice at 0.5 µL of 80 µM RNP per injected eye (40 pmol). Eyes were collected on day 14, and retinal sections were immunostained for Sox9. Müller glia reporter editing was quantified as the fraction of Sox9-positive cells that were tdTomato-positive. A parallel study used a *Vegfa*-targeting guide, with Illumina sequencing of whole-retina lysates.

### Intramuscular delivery

Under anesthesia as described above, Ai9 mice received bilateral gastrocnemius injections of 50 µL of 40 µM RNP per muscle or control. Formulations compared unmodified RNP with P55-or S315-containing PERCEPT, with or without 5-kDa PEG. On day 14, whole-animal fluorescence images were acquired and gastrocnemius muscles were harvested for ex vivo imaging and cryosection analysis. For endogenous myostatin editing, PERCEPT containing S315 and PEG was prepared with the myostatin guide (Extended Data Table 2) and dosed unilaterally for internal control.

### Intranasal delivery

Under anesthesia as described above, Ai9 mice received 25 µL of 20 µM RNP by intranasal administration. P55-and S315-containing formulations were compared with and without PEG. Lungs were collected on day 14 for imaging and histology, and reporter persistence was assessed up to 12 weeks after dosing. Pulmonary ABE delivery in GER10 mice was assessed by GFP reporter activation.

Lung sections were co-stained with rabbit anti-α-tubulin (Abcam, ab52866; clone EP1332Y), goat anti-p63/TP73L (R&D Systems, AF1916) and DAPI to assess airway cell identity. Sections were incubated overnight at 4°C with α-tubulin antibody at 1:300 and p63 antibody at 1:100 from a 1 mg/mL stock (10 µg/mL). Donkey anti-rabbit Alexa Fluor 488 and donkey anti-goat Alexa Fluor 647 were applied for 1 h at room temperature.

### Whole-animal and tissue fluorescence imaging

Whole-animal and excised-tissue epifluorescence imaging was used to assess muscle and pulmonary reporter activation; harvested gastrocnemius muscles were also imaged by IVIS at the Barker core facility, UC Berkeley. IVIS fluorescence was background-subtracted before comparison between groups.

### Targeted amplicon sequencing

Genomic DNA was extracted from the tissues specified above using the AllPrep DNA/RNA/Protein Mini Kit (QIAGEN, 80004). Target loci were amplified by PCR using the primers in Extended Data Table 3, and amplicons were submitted to Quintara Biosciences for sequencing. Reads were analyzed using CRISPResso2 version 2.3.3.^54,55^

### RNA sequencing and enrichment analysis

RNA sequencing was performed by Plasmidsaurus to profile treated R6/2 striata.^56^ Differential expression was analyzed by comparing replicate means between treatment groups using limma version 3.66.0 and edgeR version 4.8.2.^57,58^ Genes were called differentially expressed at a false discovery rate of 5% and an absolute log_2_ fold change of at least 1.

Gene set enrichment analysis was performed with GSEApy version 0.9.5.^44,59^ The analysis used the MSigDB mouse hallmark collection, mh.all.v2026.1.Mm.symbols.gmt.^60–62^ Genes were ranked by the signal-to-noise ratio between the predefined phenotype classes, and gene sets containing 15–500 genes were retained. Normalized enrichment scores and significance estimates were calculated using 1,000 permutations.

### Statistics and reproducibility

Statistical analyses used GraphPad Prism versions 10.6.0 and 11.0.0. Single-factor group comparisons used ordinary one-way ANOVA followed by Tukey’s multiple-comparisons test; the inhibitor-by-concentration experiment used ordinary two-way ANOVA with Tukey’s test. Targeted sequencing comparisons used Welch’s unpaired t-test for two groups, or one-way ANOVA with Tukey’s test for more than two groups. Data are presented as mean ± s.d. P < 0.05 was considered significant, with Tukey-adjusted P values for ANOVA comparisons.

The striatal reporter analysis included four hemispheres from two mice per condition; hemispheres were subsamples of two independent animals. Endocytic inhibitor assays used triplicate wells per condition. Muscle IVIS quantification used three separate acquisitions of harvested tissue. Myostatin sequencing included two gastrocnemius muscles per group. Each point in the lung durability analysis represented a different animal.

## Author contributions

C.B. conceived and led the study and directed material development and experimental work.

A.R. prepared materials for the lung, skeletal muscle and adenine base editor studies and contributed to experimental design for the lung and skeletal muscle studies. M.K. performed the majority of the *in vivo* experiments, including dosing and tissue histology. R.H. performed *in vitro* assays, with assistance from H.H., who also prepared materials for adenine base editor development. B.L. performed retinal administration, and J.F. advised on intravitreal delivery.

B.D.M contributed to the development of intravitreal RNP delivery. S.Z. and N.M. conceived the linker, which S.Z. synthesized. K.A. prepared the proteins. R.S. contributed to the nanogold experiments and, together with N.M. and R.W., helped develop the linker conjugation strategy.

C.G. performed next-generation sequencing. D.A. contributed computational methods for RNA-sequencing analysis. V.V.L. performed the R6/2 surgeries at Ohio State University. R.W. contributed to study conceptualization and experimental design, experimental preparation and *in vivo* studies. He supervised the research, guided interpretation of the results and contributed to manuscript preparation.

## Data & materials availability

The MATLAB script for striatal fluorescence-area quantification and its Python implementation are available at https://github.com/chrismbaehr/neuro-fluor-quant.^63^ The repository includes installation instructions, dependencies and analysis configuration.

## Acknowledgments

This work was supported by NIH Common Fund grant U19NS132303, Cystic Fibrosis Foundation grant URNOV19XX0, and by the Heritage Medical Research Institute. We acknowledge the laboratories of Deniz Dalkara and Vania Broccoli for helpful discussions and preliminary studies related to retinal delivery. We thank the UC Berkeley DNA Sequencing Facility and The IGI Next Generation Sequencing Core (NGS Core) for their support. Sarah Pyle provided assistance with graphical elements of the manuscript.

## Extended Data

**Extended Data Table 1.** Peptide sequences. Sequences are written from N to C terminus. Peptides carried a C-terminal PEG6–Cys(Tz) extension, except T1, which carried an N-terminal Cys(Tz)–PEG6 extension. The 13 peptides in Figure 1h are listed in figure order, followed by S315 used in Figure 6. The terminal extensions shown here specify the peptides used in this study.

| Peptide | Sequence with conjugation handle | Original peptide paper |
| --- | --- | --- |
| P55-Tz | KLFEAIEGFIENGWEGMIDLWNGYGRKKRRQRR–PEG6–Cys(Tz) | Sahu et al. (2025)<sup>23</sup> |
| TET1 / T1 | Cys(Tz)–PEG6–HLNILSTLWKYR | Liu et al. (2005)<sup>26</sup> |
| A22p | HTPGNSNKWKHLQENKKGRPRR–PEG6–Cys(Tz) | Shin et al. (2014)<sup>27</sup> |
| RVG variant | YTIWMPENPRPGTP–Abu–DIFTNSRGKRASN–PEG6–Cys(Tz) | Kumar et al. (2007)<sup>28</sup><br>RVG29 parent peptide. |
| KLPGWSG | KLPGWSG–PEG6–Cys(Tz) | Caprini et al. (2013)<sup>29</sup> |
| Angiopep2 | TFFYGGSRGKRNNFKTEEY–PEG6–Cys(Tz) | Demeule et al. (2008)<sup>30</sup> |
| C3 | ASKKPKRNIKA–PEG6–Cys(Tz) | Rønn et al. (1999)<sup>31</sup> |
| T7 | HAIYPRH–PEG6–Cys(Tz) | Lee et al. (2001)<sup>32</sup> |
| DRG1 | SPGARAF–PEG6–Cys(Tz) | Oi et al. (2008)<sup>33</sup> |
| DRG2 | DGPWRKM–PEG6–Cys(Tz) | Oi et al. (2008)<sup>33</sup> |
| K9_04-Tz | cyclo(E*–PEG4–LATTSQNKPA–PEG4–K*)–PEG6–Cys(Tz) | Öztürk et al. (2021)<sup>34</sup><br>Original AAV insert. |
| K9_12-Tz | cyclo(E*–LAPDSTTRSA–PEG4–K*)–PEG6–Cys(Tz) | Öztürk et al. (2021)<sup>34</sup><br>Original AAV insert. |
| K9_16-Tz | cyclo(E*–PEG4–PAPQDTTKKA–PEG4–K*)–PEG6–Cys(Tz) | Öztürk et al. (2021)<sup>34</sup><br>Original AAV insert. |
| S315-Tz | KLKLAKALAKALKLLGGSGGGSQAKAQAKQAK–PEG6–Cys(Tz) | Kulhankova et al. (2023)<sup>47</sup> |

The papers in the right-hand column report the original peptide cores or parent motifs. These citations do not describe the terminal conjugation handles used here. For RVG, the citation identifies the RVG29 parent; for the K9 peptides, it identifies the AAV capsid inserts from which the displayed cyclic scaffolds were derived.
Cys(Tz), tetrazine-functionalized cysteine; Abu, aminobutyric acid; PEG<sub>n</sub>, n ethylene glycol units; \*, Glu and Lys residues joined through a side-chain lactam bond. PEG4 segments within the K9 cyclic scaffolds are distinct from the terminal PEG6 extensions.
RVG29 reference: Kumar et al.<sup>28</sup>. The listed RVG core contains 28 residues, counting Abu as one residue, before addition of PEG6–Cys(Tz).

**Extended Data Table 2.**
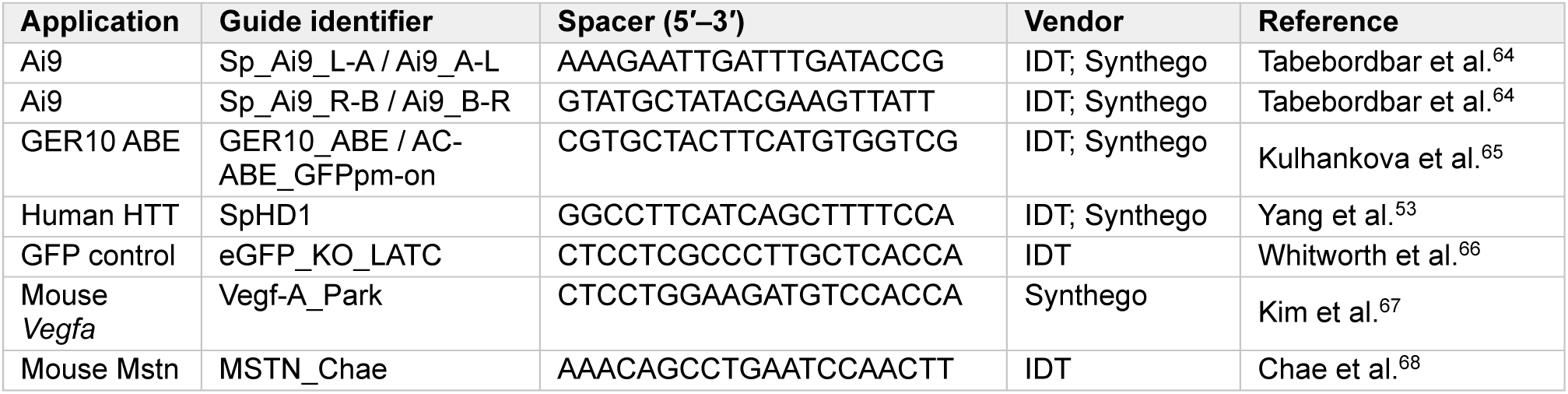
Guide RNA sequences. Spacer sequences are written 5′–3′ in the DNA alphabet (T corresponds to U in RNA). PAMs are not part of the spacer. References indicate the earliest verified publication of each guide sequence.

**Extended Data Table 3.**
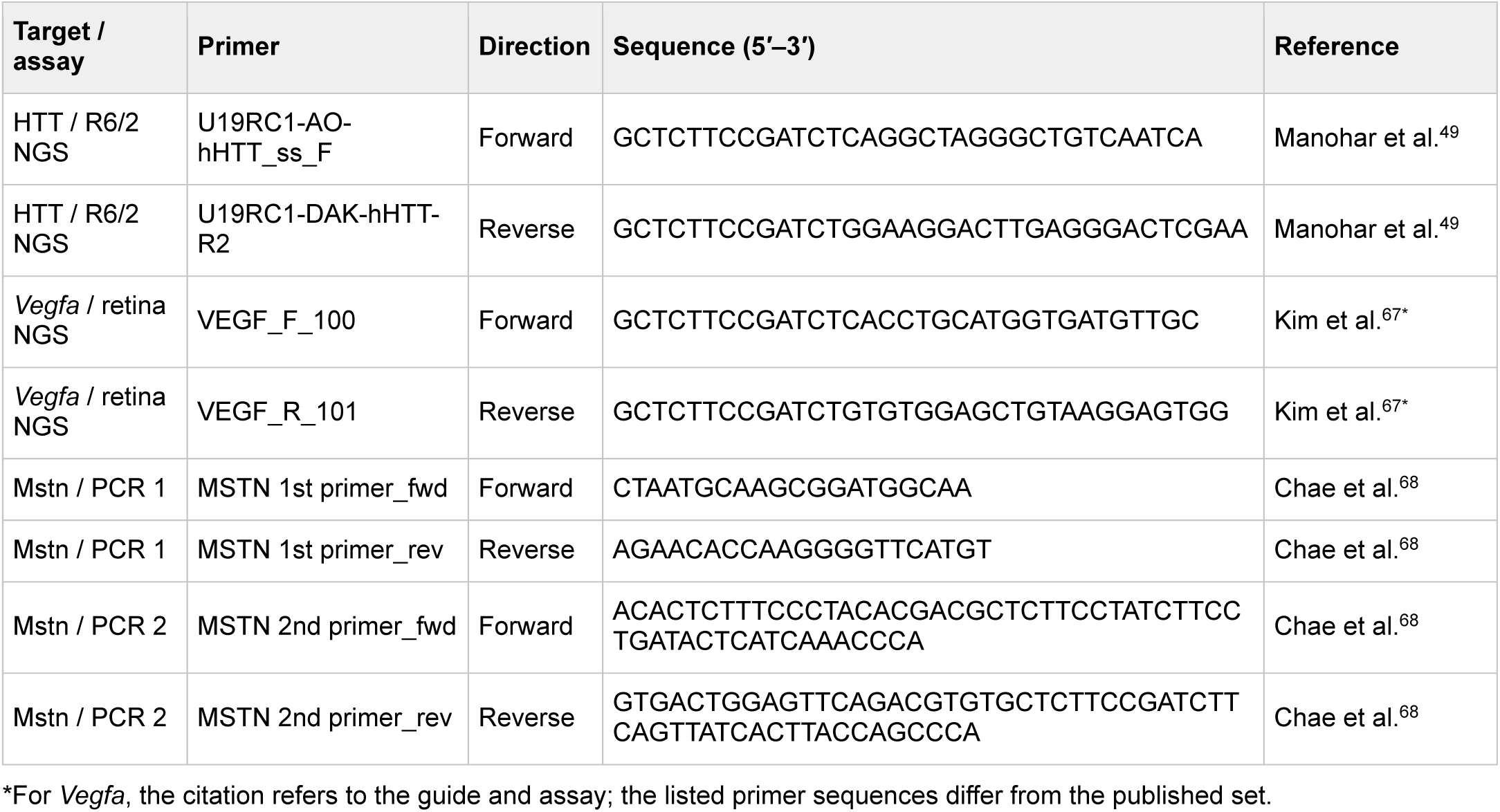
Primer sequences. Full oligonucleotide sequences are written 5′–3′, including sequencing adapters where present.

## Extended Data Figures

**Extended Data Fig. 1.**
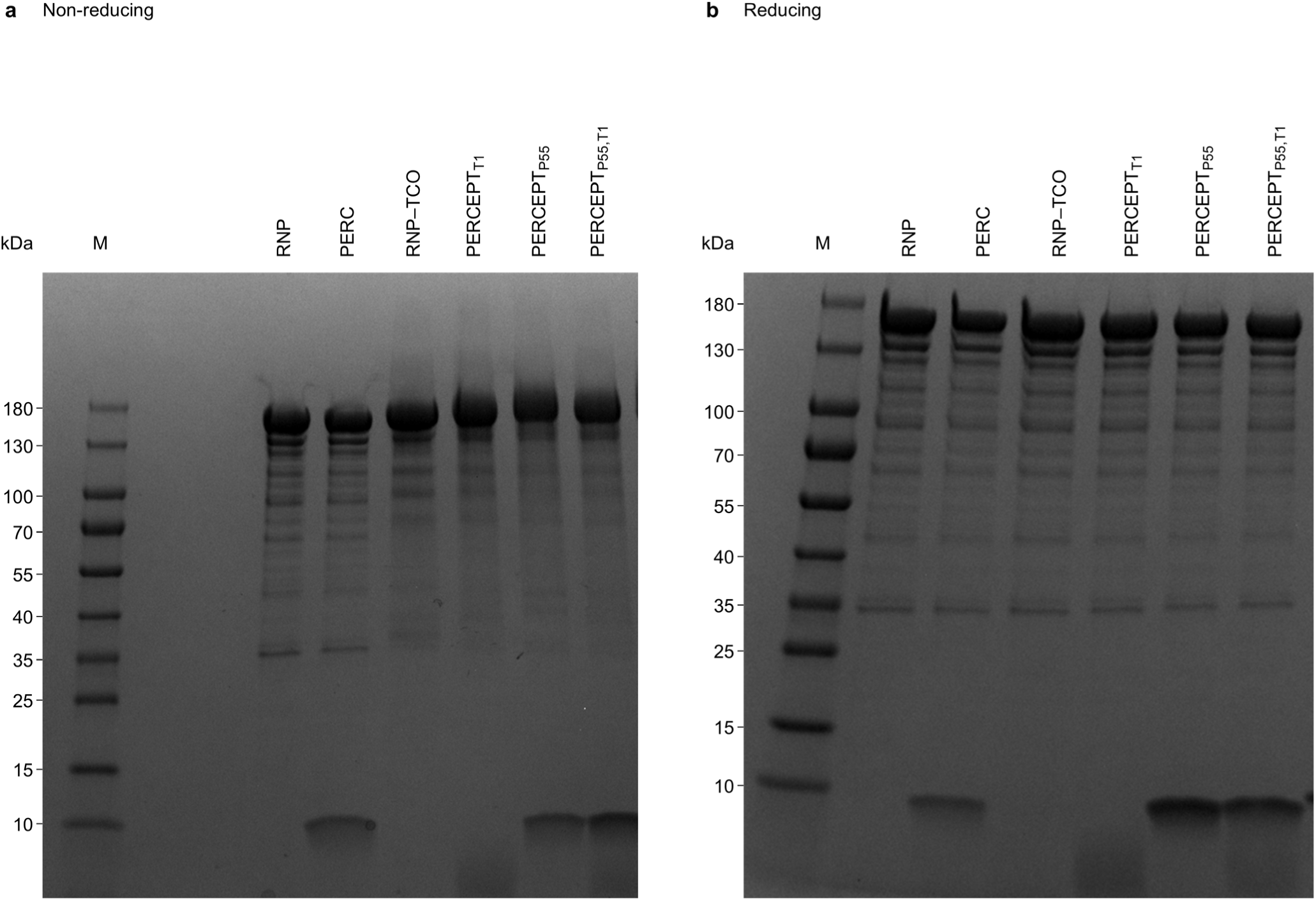
Electrophoretic analysis of RNP formulations under reducing and non-reducing conditions. Denaturing SDS–PAGE of unconjugated SpCas9 ribonucleoprotein (RNP), peptide-admixed RNP (PERC), linker-modified RNP (PERCEPT_TCO_, labelled RNP–TCO), and PERCEPT bearing T1, P55 or both peptides (PERCEPT_T1_, PERCEPT_P55_ and PERCEPT_P55,T1_, respectively). **a**, Non-reducing conditions. **b**, Reducing conditions. Reduced samples contained 2.5% β-mercaptoethanol and were heated at 98°C for 5 min. Samples were resolved on 4–20% Tris gels at 250 V for 23 min. M, PageRuler protein ladder (10–180 kDa); PERC, peptide-enabled RNP CRISPR; T1, TET1 targeting peptide; TCO, trans-cyclooctene.

**Extended Data Fig. 2.**
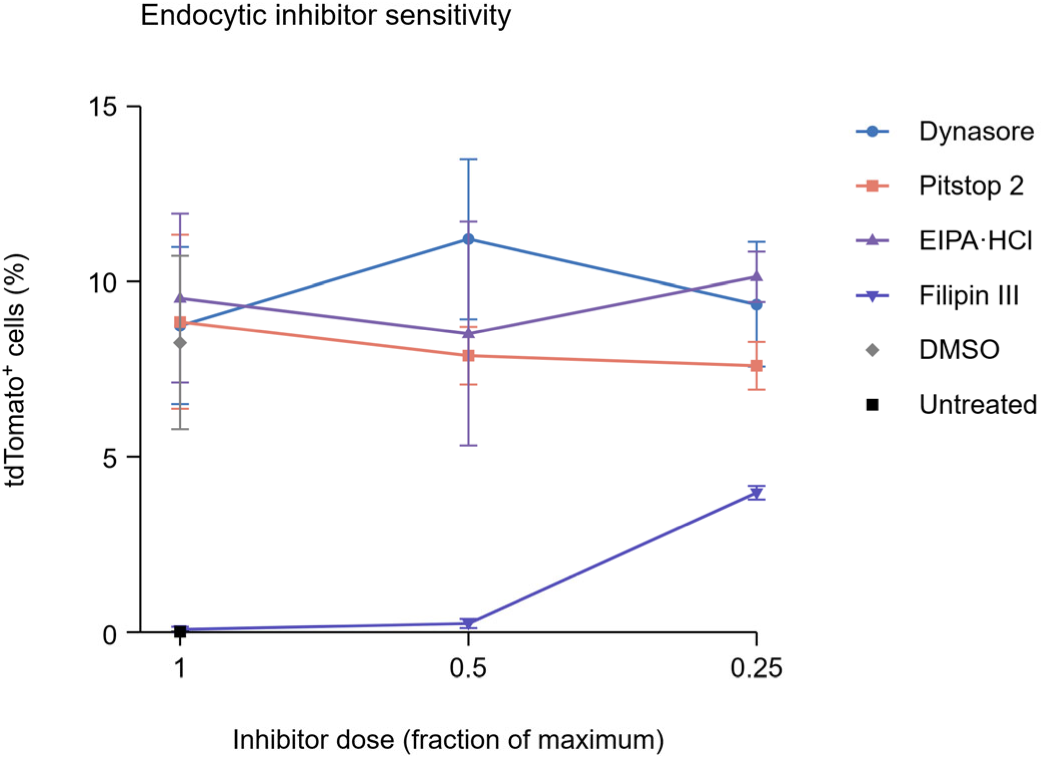
Endocytic inhibitor sensitivity of PERCEPT-mediated editing in neural progenitor cells. Flow-cytometric quantification of tdTomato reporter activation in Ai9 neural progenitor cells treated with PERCEPT ribonucleoprotein (RNP) in the presence of the indicated endocytic inhibitors. Cells received 40 pmol PERCEPT RNP during a 90-min inhibitor exposure. Inhibitor concentrations were 1×, 0.5× or 0.25× the maximum tested concentration: 30 µM Dynasore, 60 µM Pitstop 2, 200 µM EIPA hydrochloride or 4 µg ml^−1^ Filipin III. DMSO and untreated cells served as controls; control values are plotted at 1× for reference. Symbols show means and error bars denote s.d. (*n* = 3 wells per inhibitor–concentration combination; *n* = 6 and *n* = 4 observations for DMSO and untreated controls, respectively).

**Extended Data Fig. 3.**
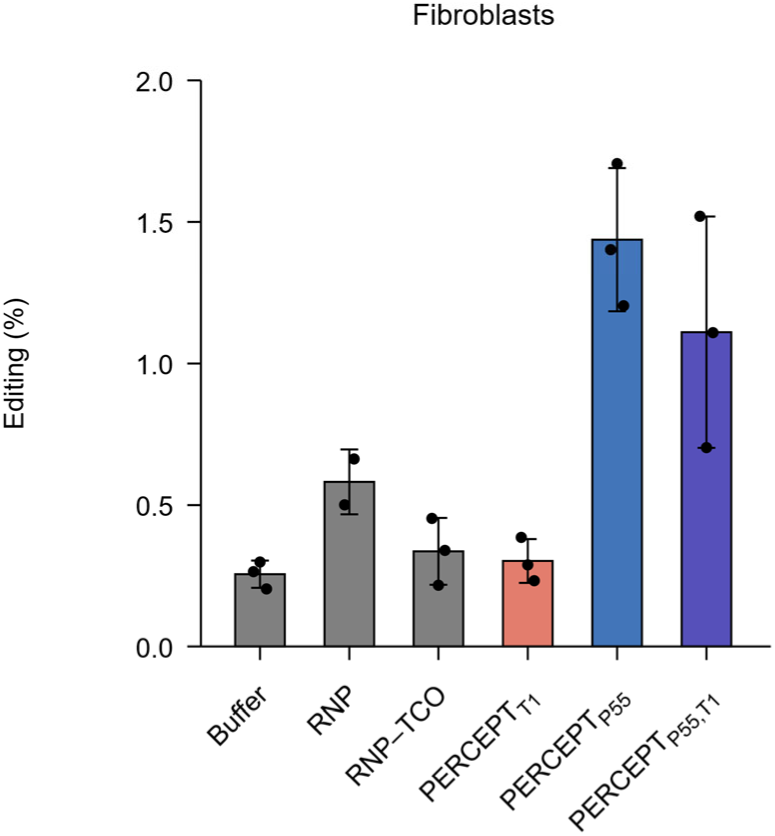
Editing activity of PERCEPT formulations in fibroblasts. Flow-cytometric quantification of editing in Ai9 fibroblasts treated with buffer, unconjugated ribonucleoprotein (RNP), linker-modified RNP (RNP–TCO), or PERCEPT bearing T1, P55 or both peptides (PERCEPT_T1_, PERCEPT_P55_ and PERCEPT_P55,T1_, respectively). T1 denotes the TET1 targeting peptide; TCO, trans-cyclooctene. Points show individual observations and bars show mean ± s.d. (*n* = 3 observations per group, except unconjugated RNP, *n* = 2).

**Extended Data Fig. 4.**
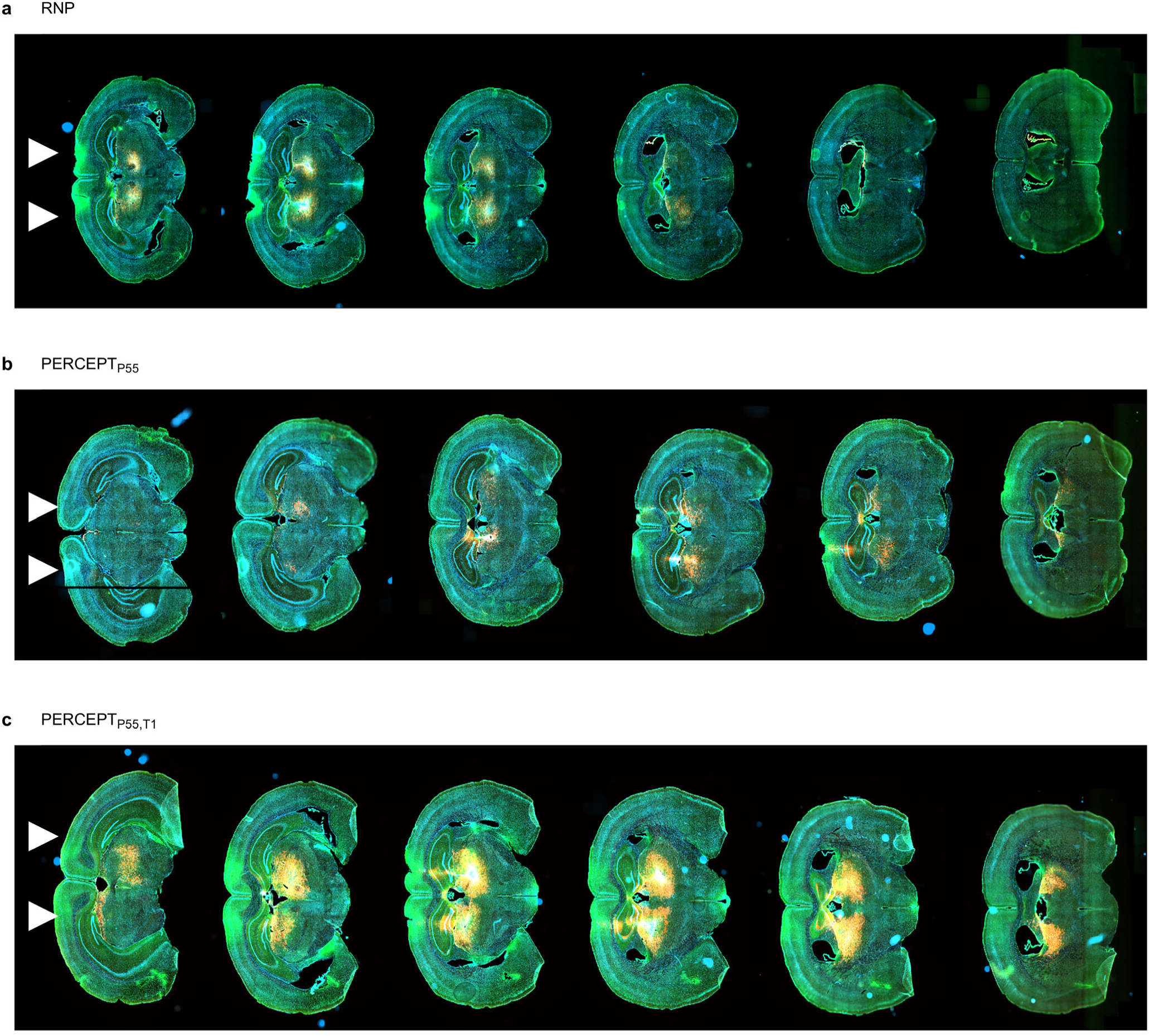
Spatial distribution of reporter activation after thalamic PERCEPT delivery. Coronal brain sections from Ai9 reporter mice after thalamic convection-enhanced delivery of Cas9 ribonucleoprotein (RNP) formulations (2 µl at 40 µM). Brains were collected 14 days after administration. **a**, Unconjugated RNP. **b**, PERCEPT_P55_. **c**, PERCEPT_P55,T1_. Six sections are shown for each formulation. White arrowheads indicate the approximate injection sites. T1 denotes the TET1 targeting peptide.

**Extended Data Fig. 5.**
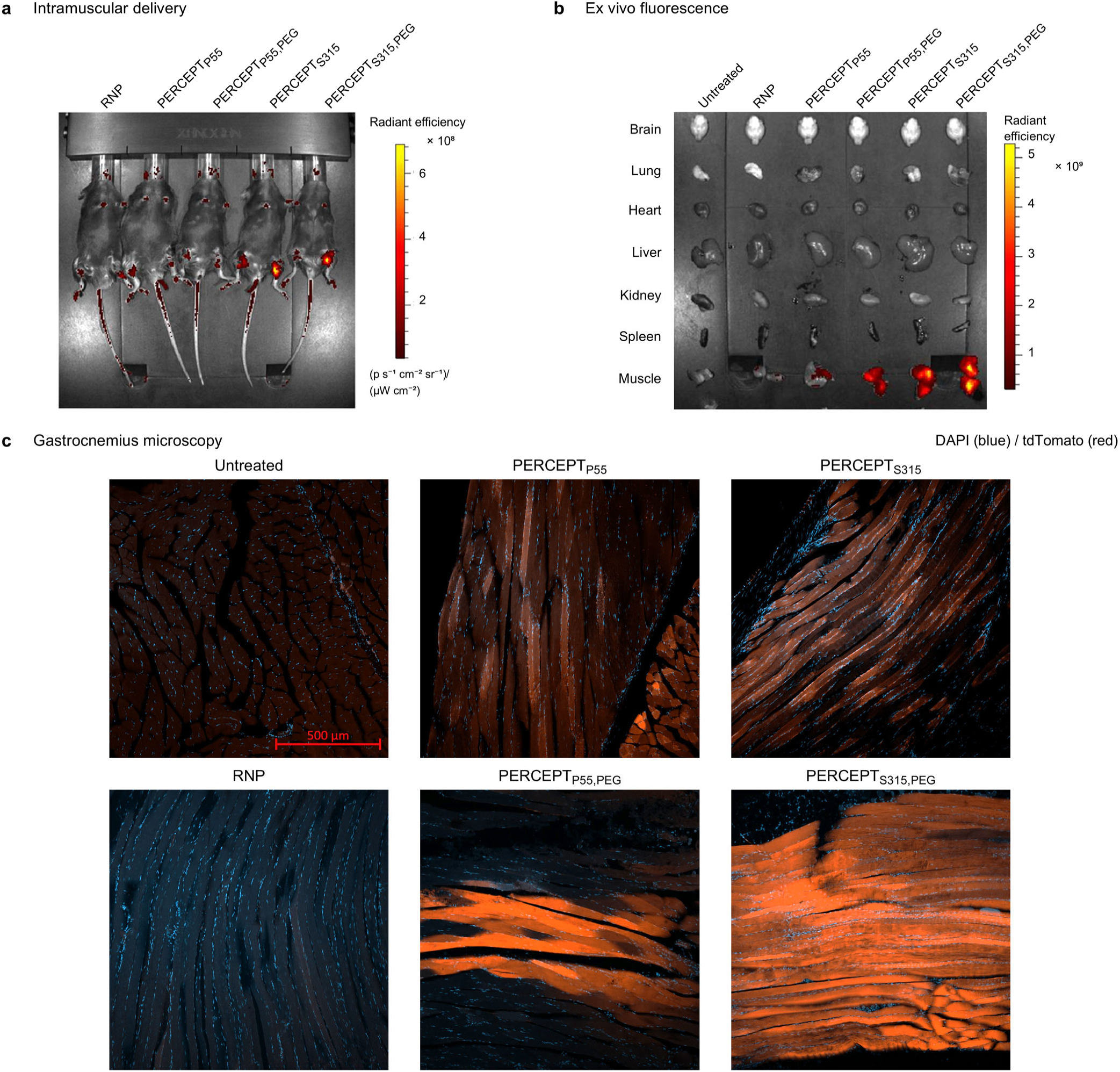
Reporter activation after intramuscular PERCEPT delivery. Ai9 reporter mice received bilateral gastrocnemius injections of 50 µl of 40 µM ribonucleoprotein (RNP) per muscle. Formulations comprised unconjugated RNP or PERCEPT bearing P55 or S315, with or without 5-kDa polyethylene glycol (PEG), as indicated. **a**, Whole-animal fluorescence images acquired 14 days after administration. **b**, Ex vivo fluorescence images of the indicated organs and tissues. Gastrocnemius muscles were harvested on day 14. **c**, Gastrocnemius cryosections showing DAPI (blue) and tdTomato (red), with untreated and unconjugated RNP controls. PERCEPT_P55_ and PERCEPT_S315_ are shown above their corresponding PEG-containing formulations, PERCEPT_P55,PEG_ and PERCEPT_S315,PEG_. Scale bar in the untreated microscopy image, 500 µm. Radiant efficiency is expressed as (photons s^−1^ cm^−2^ sr^−1^)/(µW cm^−2^); **a** and **b** use different color scales.

**Extended Data Fig. 6.**
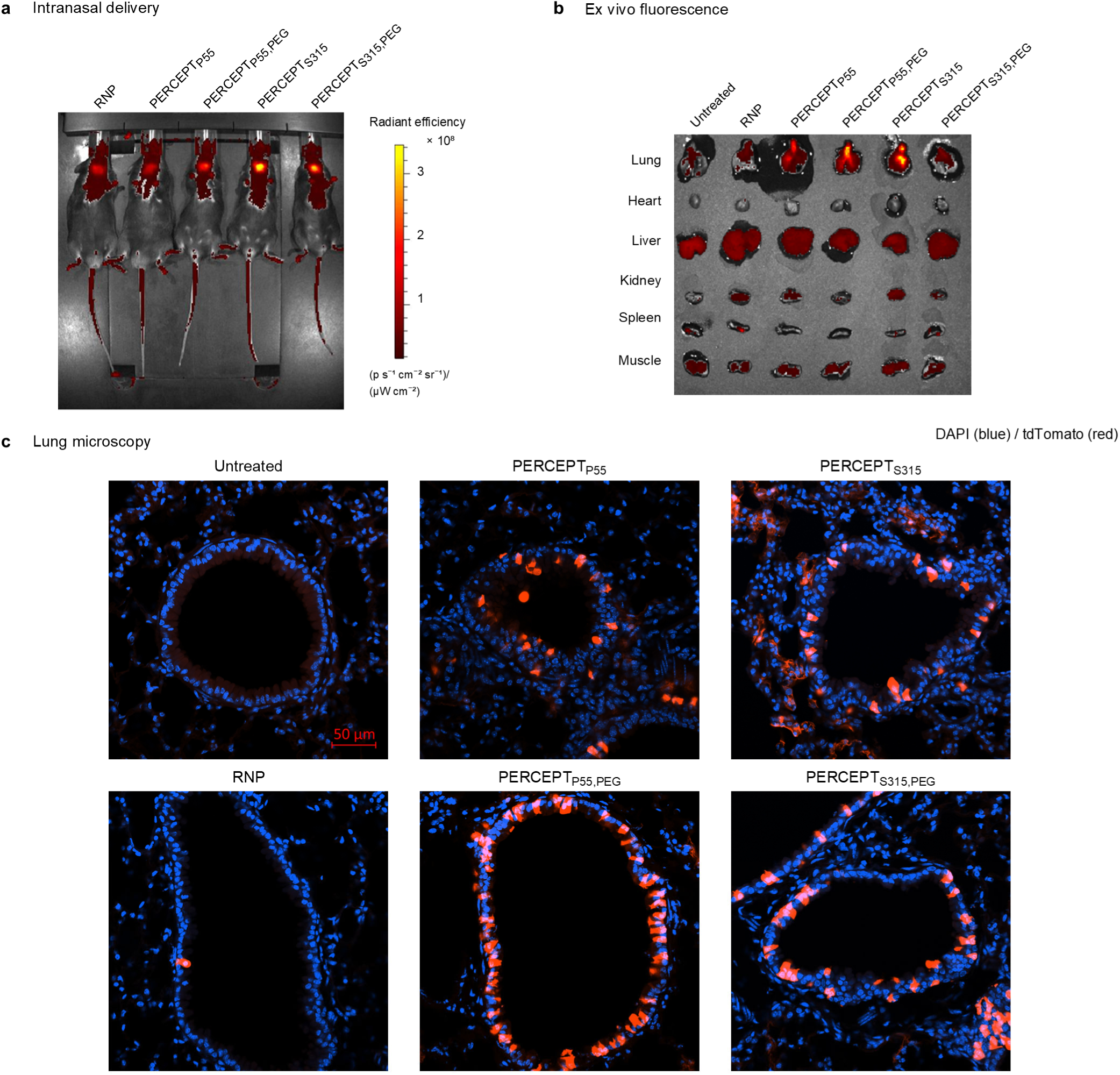
Pulmonary reporter activation after intranasal PERCEPT delivery. Ai9 reporter mice received 25 µl of 20 µM ribonucleoprotein (RNP) by intranasal administration. Formulations comprised unconjugated RNP or PERCEPT bearing P55 or S315, with or without 5-kDa polyethylene glycol (PEG), as indicated. **a**, Whole-animal fluorescence images following administration of the indicated formulations. **b**, Ex vivo fluorescence images of the indicated organs and tissues. Lungs were harvested on day 14. **c**, Lung sections showing DAPI (blue) and tdTomato (red), with untreated and unconjugated RNP controls. PERCEPT_P55_ and PERCEPT_S315_ are shown above their corresponding PEG-containing formulations, PERCEPT_P55,PEG_ and PERCEPT_S315,PEG_. Scale bar in the untreated microscopy image, 50 µm. Radiant efficiency in **a** is expressed as (photons s^−1^ cm^−2^ sr^−1^)/(µW cm^−2^).

**Extended Data Fig. 7.**
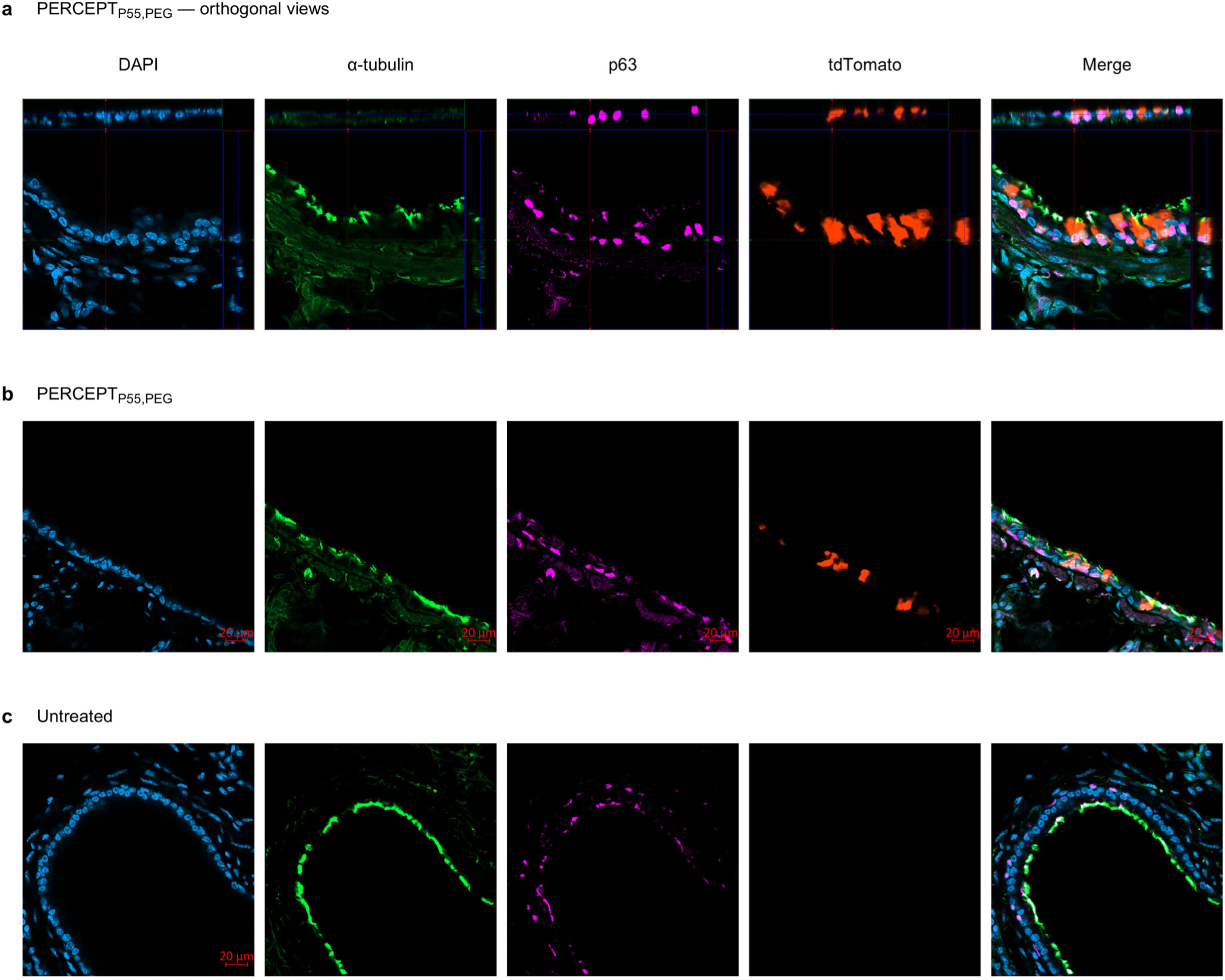
Localization of pulmonary reporter activation after PERCEPT delivery. Confocal images of lung sections from PERCEPT_P55,PEG_-treated and untreated Ai9 reporter mice. Treated mice received 25 µl of 20 µM ribonucleoprotein by intranasal administration, and lungs were collected 14 days later. **a,b**, Treated samples, shown as orthogonal views (**a**) and fluorescence images (**b**). **c**, Untreated control. Columns show DAPI (blue), α-tubulin (green), p63 (magenta), tdTomato (red) and merged channels. Scale bars, 20 µm where shown. PEG, 5-kDa polyethylene glycol.

**Extended Data Fig. 8.**
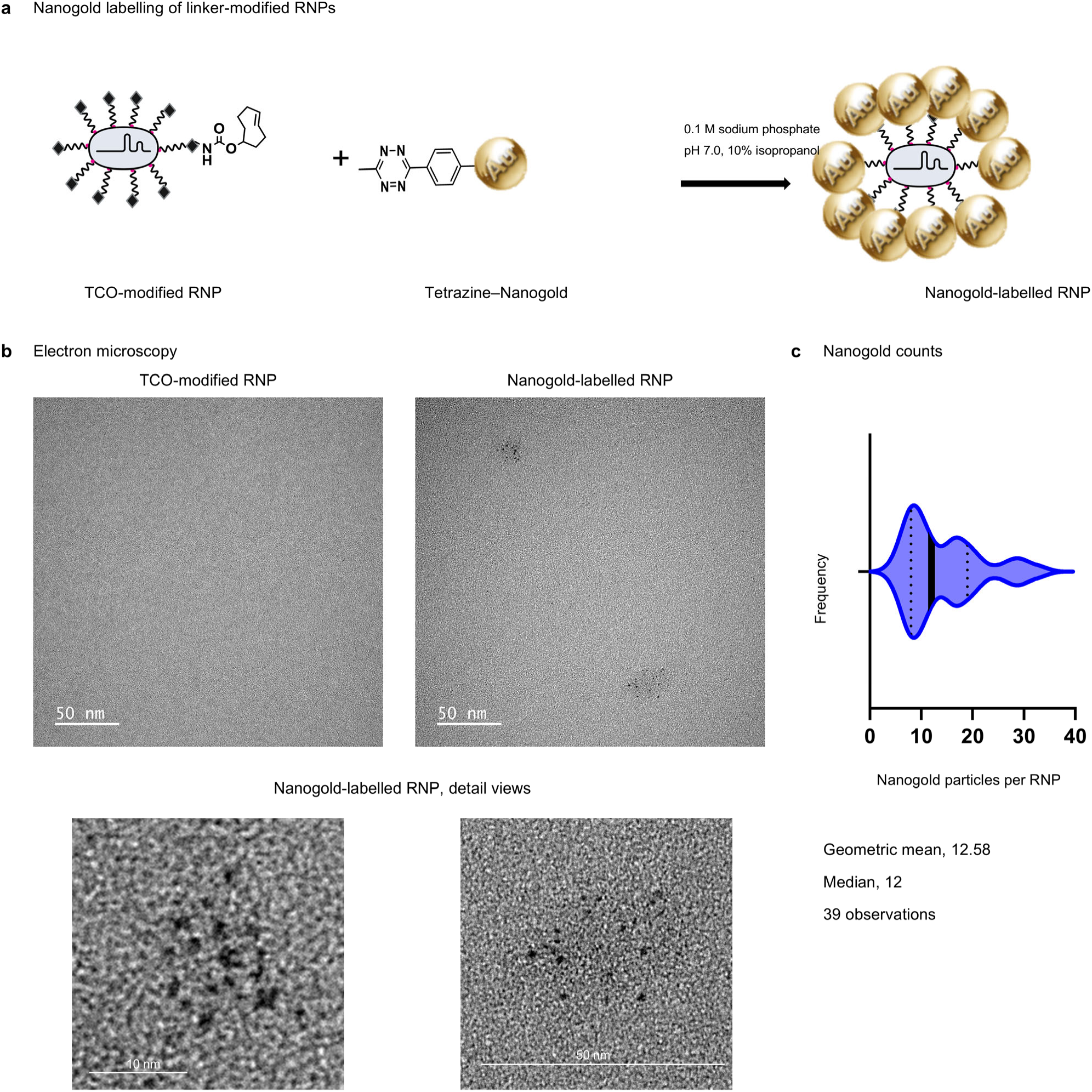
Nanogold labelling of linker-modified RNPs. **a**, Schematic of the reaction between trans-cyclooctene (TCO)-modified ribonucleoprotein (RNP) and tetrazine-functionalized Nanogold in 0.1 M sodium phosphate, pH 7.0, with 10% isopropanol. **b**, Electron micrographs of TCO-modified RNP and Nanogold-labelled RNP, with two enlarged views of labelled RNP. Scale bars, as indicated. **c**, Frequency distribution of the number of Nanogold particles per RNP (*n* = 39 observations). The geometric mean is 12.58 particles per RNP and the median is 12.

**Extended Data Fig. 9.**
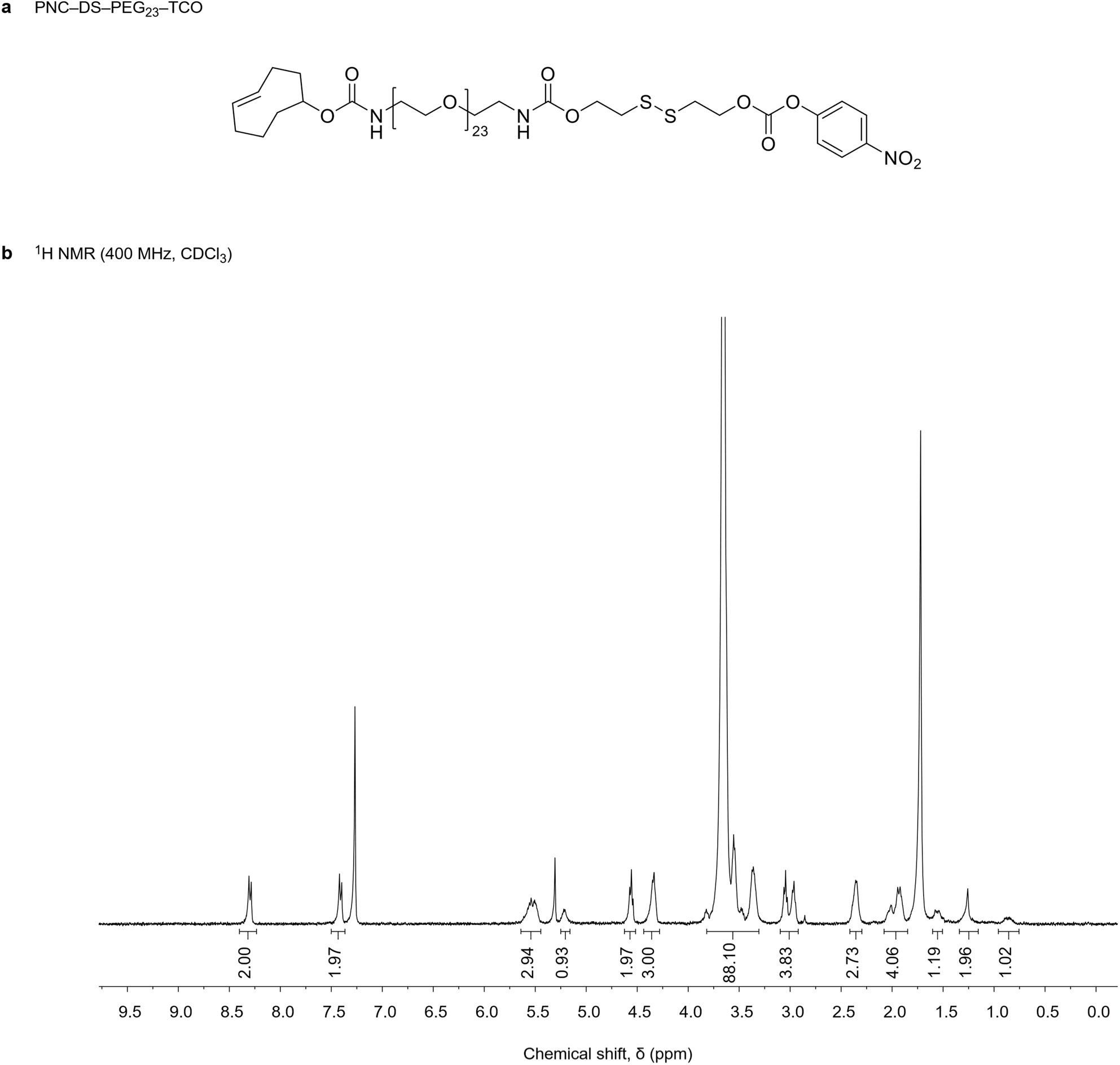
Proton NMR characterization of the PERCEPT linker. **a**, Chemical structure of PNC–DS–PEG_23_–TCO, comprising an activated para-nitrophenyl carbonate, a reduction-sensitive disulfide, a polyethylene glycol spacer containing 23 ethylene glycol units and a trans-cyclooctene handle. **b**, ^1^H NMR spectrum recorded at 400 MHz in CDCl_3_ using a Bruker AVANCE 400 spectrometer. The sample contained 2 mg linker in 600 µl solvent. Integration values are shown beneath the corresponding spectral regions.

**Extended Data Fig. 10.**
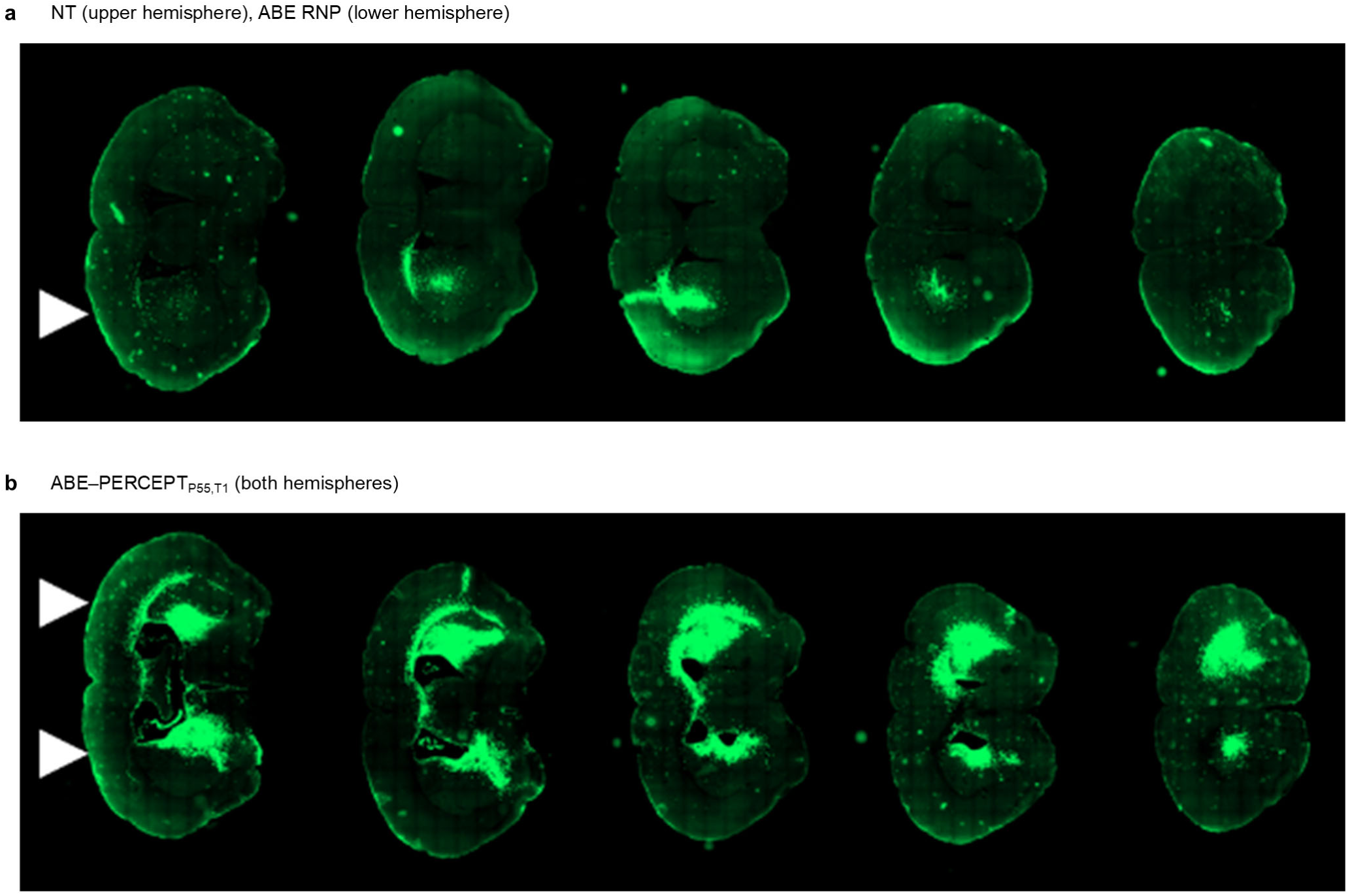
GFP reporter activation after striatal delivery of ABE RNP formulations. Coronal brain sections from GER10 reporter mice, the same mice as fig. 3e, showing GFP fluorescence (green). **a**, Untreated (NT; upper hemispheres) and unconjugated adenine base editor (ABE) ribonucleoprotein (RNP; lower hemispheres). **b**, ABE–PERCEPT_P55,T1_ in both hemispheres. Treated hemispheres received convection-enhanced delivery of 5 µl of 40 µM RNP into the striatum, and brains were collected 14 days after administration, as described in Fig. 3. Five sections are shown for each hemisphere. White arrowheads indicate the approximate injection sites.

**Extended Data Fig. 11.**
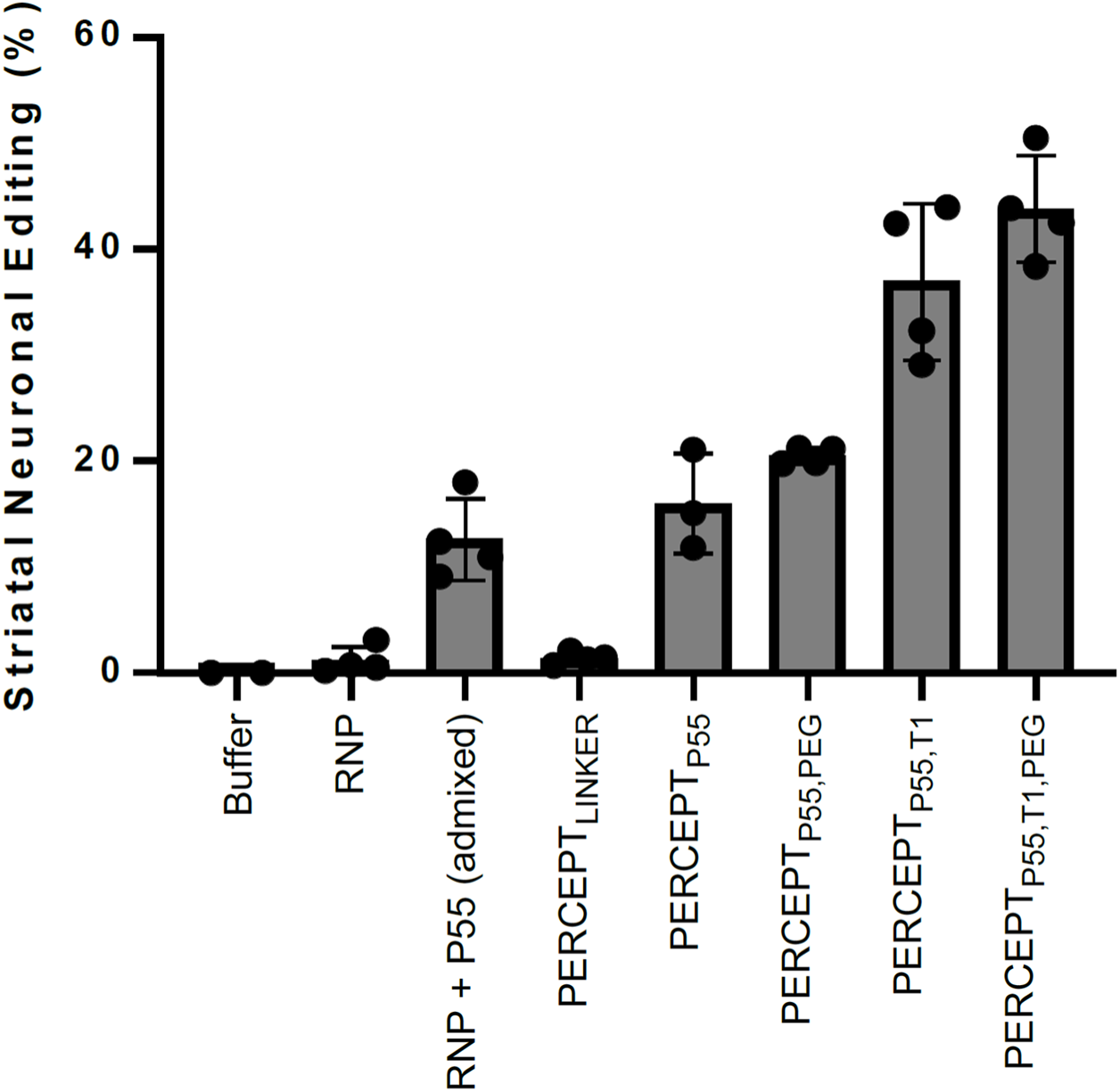
Estimate of total neuronal editing in Ai9 striatum. An estimate of the overall editing efficiency in the Ai9 striatum obtained by multiplying the values from Figures 1d & 1e (edited striatal volume & editing efficiency within that volume).

